# DNA methylation affects pre-mRNA transcriptional initiation and processing in Arabidopsis

**DOI:** 10.1101/2021.04.29.441938

**Authors:** Qiuhui Li, Shengjie Chen, Amy Wing-Sze Leung, Yaqin Liu, Yan Xin, Li Zhang, Hon-Ming Lam, Ruibang Luo, Shoudong Zhang

**Affiliations:** Department of Computer Science, The University of Hong Kong, Pokfulam, Hong Kong Special Administrative Region; School of Life Sciences, The Chinese University of Hong Kong, Shatin, Hong Kong Special Administrative Region; Center for Soybean Research of the State Key Laboratory of Agrobiotechnology, The Chinese University of Hong Kong, Shatin, Hong Kong Special Administrative Region

**Keywords:** DNA methylation, Oxford Nanopore Technology Direct RNA sequencing (ONT DRS), bisulfite sequencing, *met1-3* mutant, pre-mRNA processing, intron retention, constitutive heterochromatin, long-read sequencing, poly(A) tail

## Abstract

**Background:** DNA methylation may regulate pre-mRNA transcriptional initiation and processing, thus affecting gene expression. Unlike animal cells, plants, especially *Arabidopsis thaliana*, have relatively low DNA methylation levels, limiting our ability to observe any correlation between DNA methylation and pre-mRNA processing using typical short-read sequencing. However, with newly developed long-read sequencing technologies, such as Oxford Nanopore Technology Direct RNA sequencing (ONT DRS), combined with whole-genome bisulfite sequencing, we were able to precisely analyze the relationship between DNA methylation and pre-mRNA transcriptional initiation and processing using DNA methylation-related mutants.

**Results:** Using ONT DRS, we generated more than 2 million high-quality full-length long reads of native mRNA for each of the wild type Col-0 and mutants defective in DNA methylation, identifying a total of 117,474 isoforms. We found that low DNA methylation levels around splicing sites tended to prevent splicing events from occurring. The lengths of the poly(A) tail of mRNAs were positively correlated with DNA methylation. DNA methylation before transcription start sites or around transcription termination sites tended to result in gene-silencing or read-through events.

Furthermore, using ONT DRS, we identified novel transcripts that we could not have otherwise, since transcripts with intron retention and fusion transcripts containing the uncut intergenic sequence tend not to be exported to the cytoplasm. Using the *met1-3* mutant with activated constitutive heterochromatin regions, we confirmed the effects of DNA methylation on pre-mRNA processing.

**Conclusion:** The combination of ONT DRS with whole-genome bisulfite sequencing was a powerful tool for studying the effects of DNA methylation on splicing site selection and pre-mRNA processing, and therefore regulation of gene expression.

## Background

Pre-mRNA processing, including capping, splicing, polyadenylation site selection, and poly(A) tailing, occurs following transcription to produce mature mRNAs for subsequent translation (*1*), and it is linked to, and affected by, epigenetic modifications (*2*). Indeed, DNA methylation plays a role in pre-mRNA processing, particularly for splicing site selection, which has been reported in animal systems (*3-5*). DNA methylation also affects poly(A) site selection, because CCCTC-binding factor (CTCF), a transcription factor that regulates the 3-D structure of chromatin, only binds to unmethylated DNA, and avoids methylated targets at the 3’ end of the gene (*6*). Recently, nascent RNA profiling in *Arabidopsis* showed that more intron retention events could be observed with this method than with RNA-seq of mature mRNAs (*7*), since introns can still be spliced in the nucleus after transcription. A similar phenomenon was also observed in animal systems, called detained intron (*8*). In addition, detained introns and post-transcriptionally spliced introns are resistant to degradation by the Non-sense Mediated Decay (NMD) pathway because of the retention of these intron-containing transcripts in the nucleus (*7, 8*). However, the relationship between intron retention and DNA methylation around the splicing sites remains to be addressed.

A poly(A) tail is an important signature of mature mRNAs, and facilitates mRNA export to the cytoplasm for translation (*9*). The length of a poly(A) tail has also attracted attention for its implications in mRNA stability. It was found that a short poly(A) tail is a conserved feature of housekeeping genes (*10*), as confirmed by whole-transcriptomic poly(A) length sequencing (PAL SEQ) (*11*). A longer poly(A) tail may facilitate mRNA degradation via the exosome pathway, which is dependent on a poly(A) binding protein (PABPN1) and poly(A) polymerases (PAPs - specifically PAPα and PAPγ in mammalian systems) (*12*). The poly(A) tail extension in the nucleus is controlled by a poly(A) binding protein which regulates the interaction between poly(A) polymerase and cleavage, and a poly(A) specificity factor (*13*). However, the role of DNA methylation on poly(A) tail length determination remains unclear.

Typically in RNA sequencing, mRNAs are first reverse-transcribed to synthesize the first-strand complementary DNA (cDNA) which are then used as templates to amplify cDNA (*14*). The amplified cDNA can then be subjected to Sanger sequencing, next-generation sequencing, or third-generation sequencing (e.g. PacBio (*15*)] or ONT cDNA sequencing (*7*)) to determine mRNA sequences. Although the classical method helps us determine gene expression levels and understand pre-mRNA processing, its limitations prevent us from fully understanding mRNA molecular traits. For example, current commercially available reverse transcriptases may not reverse-transcribe RNA molecules longer than 12,000 nt (https://www.thermofisher.com/hk/en/home/life-science/cloning/cloning-learning-center/invitrogen-school-of-molecular-biology/rt-education/reverse-transcription-setup.html), so it cannot provide cDNAs longer than 12,000 nt for sequencing. Moreover, some modifications and secondary structures of RNA molecules may inhibit the activity and/or decrease the fidelity of reverse transcriptases (*16-18*), and thus we may not get the full-length cDNAs of these RNA molecules. In recent years, Oxford Nanopore developed a novel sequencing method, Oxford Nanopore Technology Direct RNA sequencing (ONT DRS), to directly sequence native RNA molecules, which is therefore not limited by RNA molecular lengths and can document modifications on RNA bases (*19*). ONT DRS has been well used in both animal (*20*) and plant systems (*21, 22*), resulting in many new insights into RNA molecules. Although base-calling for ONT DRS is still not as accurate as next-generation sequencing, if a well-annotated reference genome is available, the imperfect base-calling should not interfere with downstream analyses (*21*).

DNA methylation is a crucial epigenetic mark determining gene expression and heterochromatin maintenance (*23*), but there is a difference between DNA methylation patterns in mammalian and plant systems. In mammalian cells, DNA methylation occurs mainly in the CG context, while in plants, DNA methylation may occur in the CG, CHG and CHH contexts. In *A. thaliana*, DNA methyltransferase 1 (MET1) is responsible for CG methylation, so loss-of-function mutants of *MET1* will nearly eliminate CG methylation genome-wide. Chromomethylase 3 (CMT3) is mainly responsible for maintaining CHG methylation, while domain-rearranged methyltransferase 1 (DRM1), DRM2, and CMT2 are responsible for *de novo* CHH methylation (*24*). Therefore knocking out all four genes (*DRM1, DRM2, CMT2* and *CMT3*) would almost completely deplete CHG and CHH methylation in the *ddcc* mutant (*3*). In *A. thaliana*, the DREAM complex, containing TCX5 (Tesmin/TSO1-like CXC domain-containing protein 5) and TCX6, could inhibit the expressions of *MET1, CMT3*, and some other DNA methylation-related genes, such as *VIM1* (*VARIANT IN METHYLATION 1*) (*25*). Thus, the double mutant *tcx5tcx6* does not have a functional DREAM complex, resulting in increased genome-wide DNA methylation (*25*). In this study, we used these DNA methylation-related mutants in a combination of ONT DRS and whole-genome bisulfite sequencing to investigate DNA methylation effects on pre-mRNA transcription initiation and processing, and found that it played a key role in pre-mRNA transcription initiation, splicing site selection, transcription termination site selection, and poly(A) length determination. We also found that for fusion transcripts, if the transcribed intergenic regions were not spliced, these transcripts tended to remain in the nucleus, but if they were fully spliced, they would be exported to the cytoplasm. Using the activated constitutive heterochromatin in *met1-3*, we further confirmed the relationship between DNA methylation and pre-mRNA transcriptional initiation and processing we observed in the euchromatin regions.

## Results

### Oxford Nanopore Technologies Direct RNA Sequencing (ONT DRS) revealed the pattern of gene expression disruption by T-DNA

Our previous ONT DRS study indicated that ONT DRS can effectively sequence native full-length RNA molecules (*21*) and thus capture relatively accurately real-time RNA dynamics at the time of harvest of the plant materials. In this study, we extracted RNA from 12-day-old seedlings of the DNA methylation-related mutants, *met1-3* (*26*), *ddcc* (*3*) and *tcx5tcx6* (*25*) along with wild type (Col-0), to profile the relationship between DNA methylation and pre-mRNA transcription initiation and processing using ONT DRS combined with whole-genome bisulfite sequencing (BS-Seq). The ONT DRS library of each genotype contained more than 2 million full-length reads with a single flow cell (R9.4.1), except for *tcx5tcx6*, for which more than 2 million full-length reads were obtained with two flow cells. The read quality and average length were similar to our previous ONT DRS data (*21*); Additional file 1: Table S1).

Most of these DNA methylation-related mutants were produced by T-DNA insertions, except *tcx6* which was created via CRISPR-Cas9-mediated gene editing (*25*). Since our ONT DRS can sequence full-length RNA molecules, we wanted to know if we could use this technique to decipher how T-DNA insertions affected the expression of the mutated genes. We used the ONT DRS full-length reads covering the T-DNA-inserted genes to examine the transcriptional dynamics of these genes. For the *MET1* gene in *met1-3* and the *CMT3* gene in the *ddcc* quadruple mutant, the DNA sequences on both sides of the T-DNA insertion sites were well-transcribed, but we could find no reads spanning both sides of the insertion sites. It seems that the pre-insertion-site mRNAs were transcribed using the promoter of the affected gene for initiation, while those post-insertion-site mRNAs were transcribed starting from within the inserted T-DNA, and reads from both sides of the insertion sites were clearly separated by the T-DNA due to its unusually long sequence (Additional file 2: Fig. S1). For *DRM2* and *CMT2* in the *ddcc* mutant, only transcripts initiated at the original promoters of these affected genes and ending before the insertion sites could be found. No post-insertion-site transcripts were obtained (Additional file 2: Fig. S1). One explanation for this phenomenon may be that intact T-DNAs always include transcription termination sites (TTSs) in their sequences, so even though the T-DNAs may be short enough for readthrough, the transcripts that were initiated from upstream of the insertion sites will still be terminated within the T-DNA. For transcribing the sequences downstream of the insertion sites, even if sometimes the TTSs within the T-DNA sequences may be lost due to truncation during insertion into target genes (*27*), the introduced promoters within the T-DNA sequences may be used for initiating transcription. For the *DRM1* gene in the *ddcc* mutant, only one post-insertion-site read was obtained. The *DRM1* gene was not usually expressed in Col-0 in normal conditions. No read was mapped to the *TCX5* locus in the *tcx5tcx6* double mutant, although the gene contains a T-DNA insertion as verified by genotyping. All the above data confirmed that the mutants used in the study cannot produce full-length functional transcripts from the T-DNA-inserted genes.

### Whole-genome bisulfite sequencing (BS-seq) combined with ONT DRS indicated a negative correlation between DNA methylation and gene expression in constitutive heterochromatin regions

To investigate the relationship between DNA methylation and gene expression, we first evaluated the alterations in genome-wide DNA methylation patterns in the DNA methylation-related mutants, using our data from BS-seq and those downloaded from the GEO database (accession no. GSE137754 for the *tcx5tcx6* mutant). The differential DNA methylation regions (DMRs) between the mutants and wild type were identified on all five chromosomes (Fig. 1, Additional file 2: Fig. S2). The results showed that CG methylation was nearly eliminated in *met1-3* mutants since the *MET1* gene was knocked out, this being especially obvious in the centromeric regions in the CG context.

**Fig. 1.**
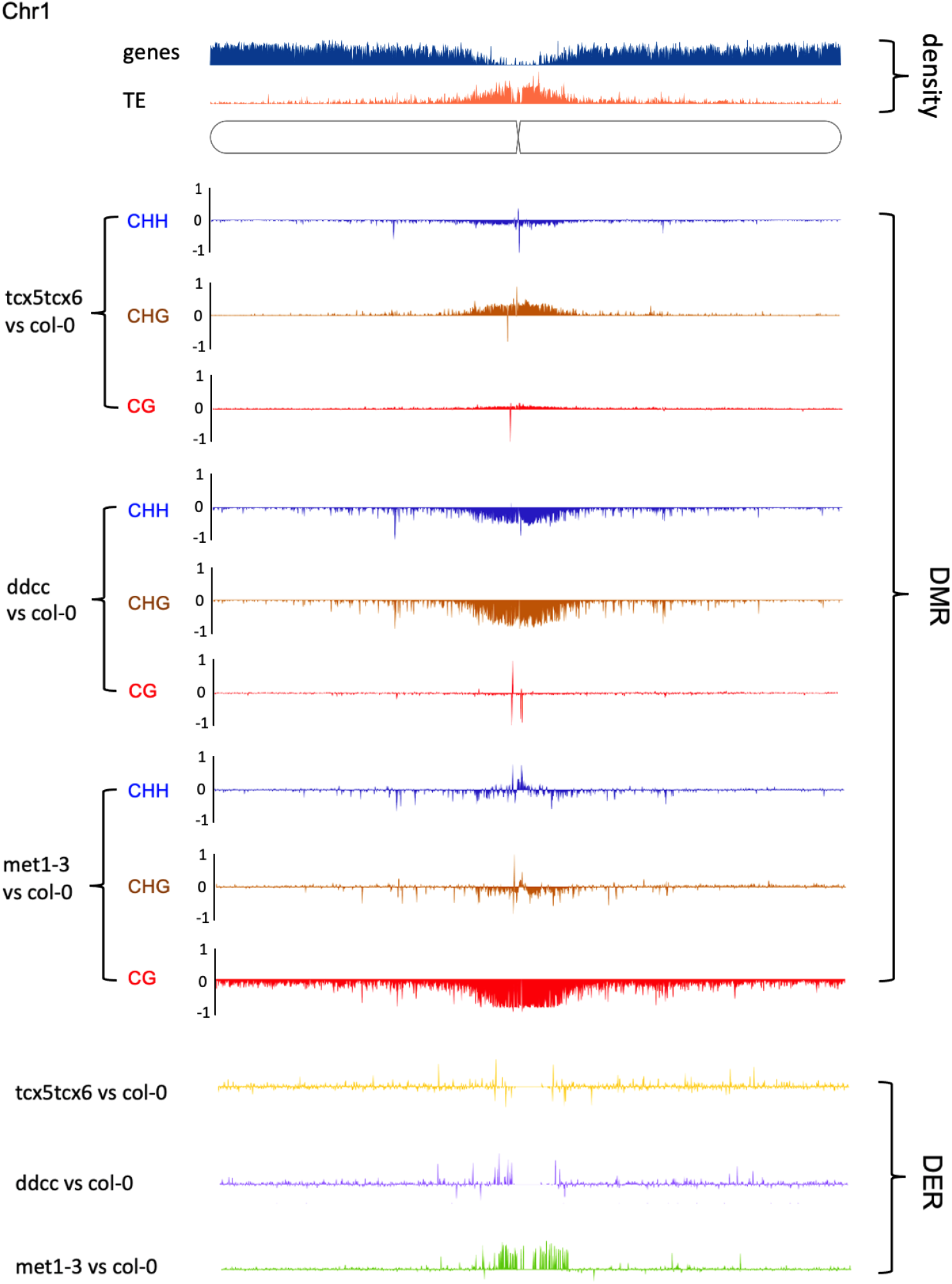
Comparative distributions of the differential expression regions (DER) and differential methylation regions (DMR) along Chromosome 1 (Chr1) between individual DNA methylation-related mutants (*met1-3*/*ddcc*/*tcx5tcx6*) and wild type (Col-0) of Arabidopsis at the CHH, CHG and CG contexts.

Effects on DNA methylation in the CHG and CHH contexts in the centromeric regions could also be observed in this mutant, but to a much lower degree (Fig. 1, Additional file 2: Fig. S2). In the *ddcc* quadruple mutant, most CHG and CHH methylations were reduced, particularly in the centromeric regions. However, DNA methylation in the CG context in the *ddcc* quadruple mutant was only slightly affected in the centromeric regions and genome-wide (Fig. 1, Additional file 2: Fig. S2). The DNA methylation patterns of *tcx5tcx6* were also analyzed using the published BS-seq data using 12-day-old seedlings (*25*), same as what was used for ONT DRS in this study. The DNA methylation in *tcx5tcx6* seedlings increased slightly in the CG and CHG contexts, and decreased slightly in the CHH context for all 5 chromosomes (Fig. 1, Additional file 2: Fig. S2). All the observed DMR patterns were as expected for these DNA methylation-related mutants.

To investigate DNA methylation effects on gene expression, we analyzed differentially expressed regions (DERs) between mutants and the wild type control (Col-0) using the ONT DRS data. After mapping the DERs to the chromosomes, we observed a striking activation in the hypomethylated centromeric regions in all five chromosomes in the *met1-3* mutant, but not in other mutants (Fig. 1, Additional file 2: Fig. S2). Although DERs are not always tightly coupled with the methylation level, down-regulated DERs generally has slightly higher DNA methylation, as seen in the *tcx5tcx6* double mutant (Fig. 1, Additional file 2: Fig. S2). Our results showed that DNA methylation, especially in the CG context, had a strong negative correlation with gene expressions in the constitutive heterochromatin regions (centromeric regions), but such a negative correlation could not be established in euchromatin regions (Fig. 1, Additional file 2: Fig. S2), since DNA methylation in the gene body and the promoter may have different effects on gene expression (*28, 29*).

### DNA methylation was positively correlated with poly(A) tail length

The poly(A) tail is crucial for mRNA stability and export to the cytoplasm (*30*). Our ONT DRS results clearly showed the difference in poly(A) tail lengths among housekeeping genes (*31*), other (non-housekeeping) genes, and transposable element (TE) genes in all four genotypes (Fig. 2). In general, in all the genotypes, the housekeeping genes had a slightly shorter poly(A) tail (with a mode of about 60 nt) than other (non-housekeeping) genes, which is consistent with recent reports of highly expressed genes tending to have a shorter poly(A) tail (*10, 11, 32*), and the actual values were similar to previous reports of the poly(A) lengths of *Arabidopsis* housekeeping genes (*11*). On the other hand, transcripts from TE regions tended to have the longest poly(A) tails in all the genotypes (Fig. 2). Although the poly(A) tail lengths varied among different transcripts, there was no significant difference in the lengths of poly(A) tails in the same transcript category among genotypes (Additional file 2: Fig. S3). Since DNA methylation in the TE regions tends to be hypermethylated, and the transcripts from these regions tend to have a longer poly(A) tail, we wanted to test whether the length of the poly(A) tail is correlated with the level of DNA methylation in the corresponding genomic regions. We first analyzed the distribution of DNA methylation levels in the genomic regions corresponding to the housekeeping genes, other genes and TE genes for all four genotypes (Additional file 2: Fig. S4). The CHG and CHH methylation data showed that housekeeping genes tended to have the lowest DNA methylation level, while the regions encoding TEs had the highest in nearly all genotypes, except in *ddcc* which had almost no CHG and CHH methylation. CG methylation of housekeeping genes was the highest among the three categories of genes, while there was no significant difference in CG methylation levels for TE and other genes across all genotypes, except *met1-3* which had next to no CG methylation at all (Additional file 2: Fig. S4). It is known that poly(A) tail formation requires poly(A) polymerases to function at the cleavage sites produced by Cleavage and Poly(A) Specificity Factor 73 (CPSF-73) (*33, 34*), and that the activity of poly(A) polymerases may be affected by DNA methylation around the TTS because the processing of the 3’ ends of pre-mRNAs is concurrent with transcription (*35*). Therefore, we compared DNA methylation levels around the TTS of the genomic regions corresponding to the three kinds of transcripts. From our results, it is clear that DNA methylation around the TTS is highest for TE genes, and lowest for housekeeping genes in all contexts, including CG, CHG and CHH, in nearly all genotypes, except for CHG and CHH methylation that were abolished in the *ddcc* quadruple mutant and CG methylation which was abolished in the *met1-3* mutant (Fig. 3). The results strongly suggested that DNA methylation was positively correlated with poly(A) tail length.

**Fig. 2.**
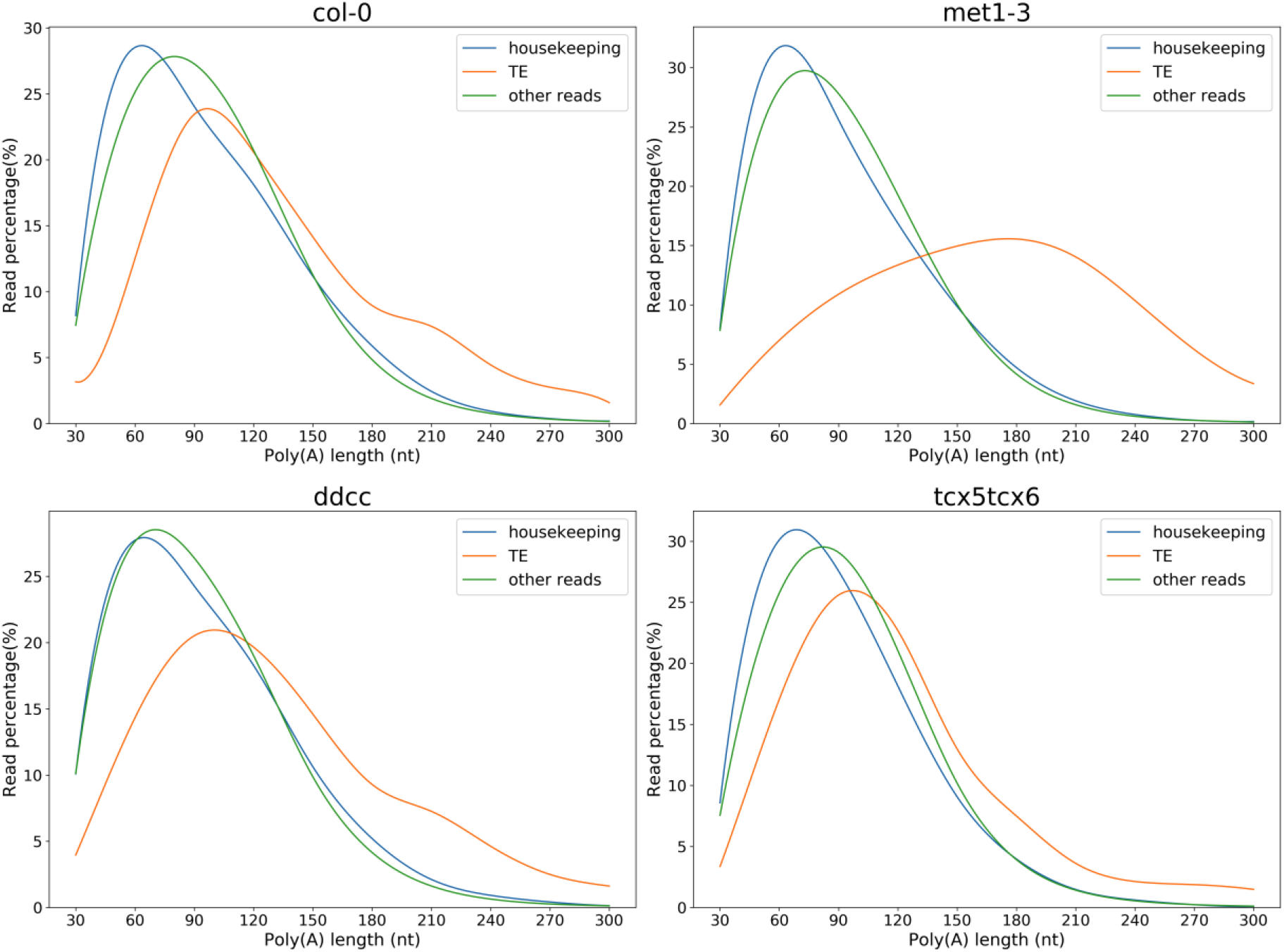
Distributions of poly(A) tail lengths of housekeeping genes, transposable element (TE) genes, and other genes in the DNA methylation-related mutants (*met1-3*/*ddcc*/*tcx5tcx6*) and wild type (Col-0) of Arabidopsis.

**Fig. 3.**
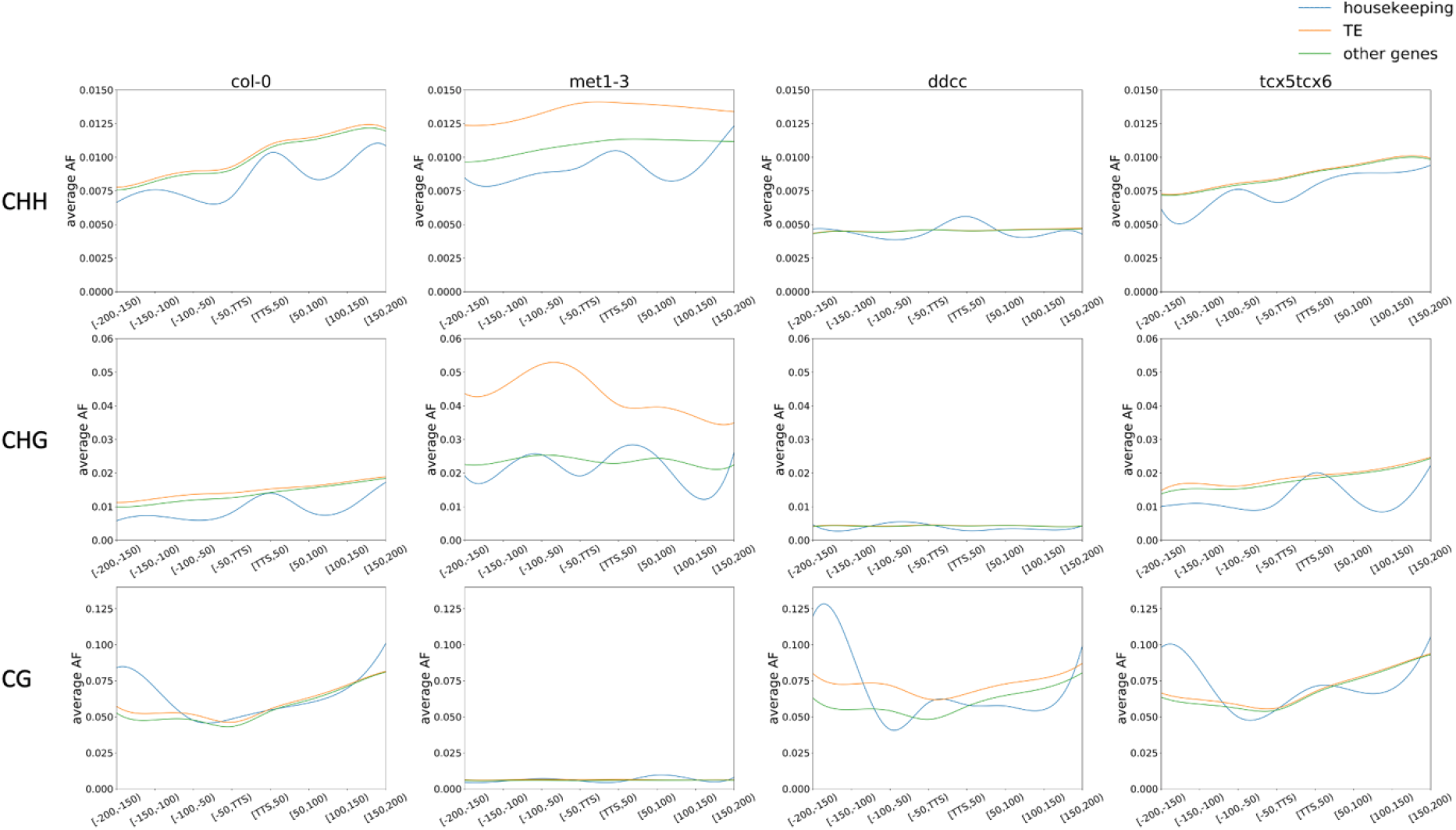
DNA methylation patterns around the transcriptional termination sites (TTSs) of housekeeping genes, TE genes and other genes in the DNA methylation-related mutants (*met1-3*/*ddcc*/*tcx5tcx6*) and wild type (Col-0) of Arabidopsis at the CHH, CHG and CG contexts.

### DNA methylation levels may determine the usage of splicing sites

We adapted the published algorithm, TrackCluster (TC) (*1, 9*), for use in analyzing RNA splicing sites in *A. thaliana* and renamed it TC-RENO (https://github.com/HKU-BAL/TC-RENO). Using TC-RENO, we identified 8,881 intron retention events, 263 extra exon events, and 729 exon-skipping events among all four genotypes. The numbers of events in each of these categories of alternative splicing that were unique to one genotype were far lower than the numbers of those that were common to all four different genotypes (Additional file 2: Figs. S5-S7). This is also true when we compared each of the mutants separately to Col-0. The unique events in Col-0 may reflect the loss of such events in the mutants, and unique events in mutants but not in Col-0 indicate a gain of such events. Generally, there was no obvious difference between the amount of loss or gain of these events in any mutants compared to the wild type, except for a slightly higher gain than loss of extra exon events and exon-skipping events in *ddcc* and *met1-3*, and a slightly higher loss than gain of extra exon and exon-skipping in *tcx5tcx6* (Additional file 2: Fig. S6, Fig. S7). For intron retention events, there was a slight decrease in gain compared to loss of unique intron retention events in all mutants (Additional file 2: Fig. S5). To understand this phenomenon, we calculated the number of donor sites and acceptor sites for each sample. Not surprisingly, most of the splicing sites (both donor and acceptor sites) belonged to splicing sites common to all four genotypes (Additional file 2: Fig. S8, Fig. S9), while *met1-3* and *tcx5tcx6* had a slightly higher number of unique splicing sites compared to Col-0 and *ddcc*.

Common events between Col-0 and mutants or among genotypes are mostly obvious events, and intron retention events are indeed the most common alternative splicing events in *A. thaliana* (*36*). Here we performed whole-genome BS-seq for Col-0 as well as *ddcc* and *met1-3* and downloaded the BS-seq data of *tcx5tcx6* conducted at the same developmental stage as the ONT DRS here using *tcx5tcx6* seedlings, to investigate the DNA methylation patterns around the splicing sites of the intron retention events at the corresponding loci and at those housekeeping and other gene loci corresponding to the fully spliced events. Surprisingly, we found that DNA methylation in the CG context around the splicing sites of intron-retained genes was distinctively lower than that around fully spliced sites of housekeeping genes and other genes (Fig. 4) in nearly all genotypes except for *met1-3*, since CG methylation in the mutant was nearly eliminated (Fig. 1, Additional file 2: Fig. S2). CHH and CHG methylation around the splicing sites (donor or acceptor) on the intron sides had slightly higher DNA methylation in intron-retained genes than those in housekeeping and other genes without intron retention events, except for the *ddcc* mutant where CHG and CHH methylation were nearly eliminated (Fig. 4). Since all the samples displayed the same pattern, we concluded that DNA methylation does play a role in intron retention, and the correlation is only with DNA methylation around the splicing sites, regardless of genotype.

**Fig. 4.**
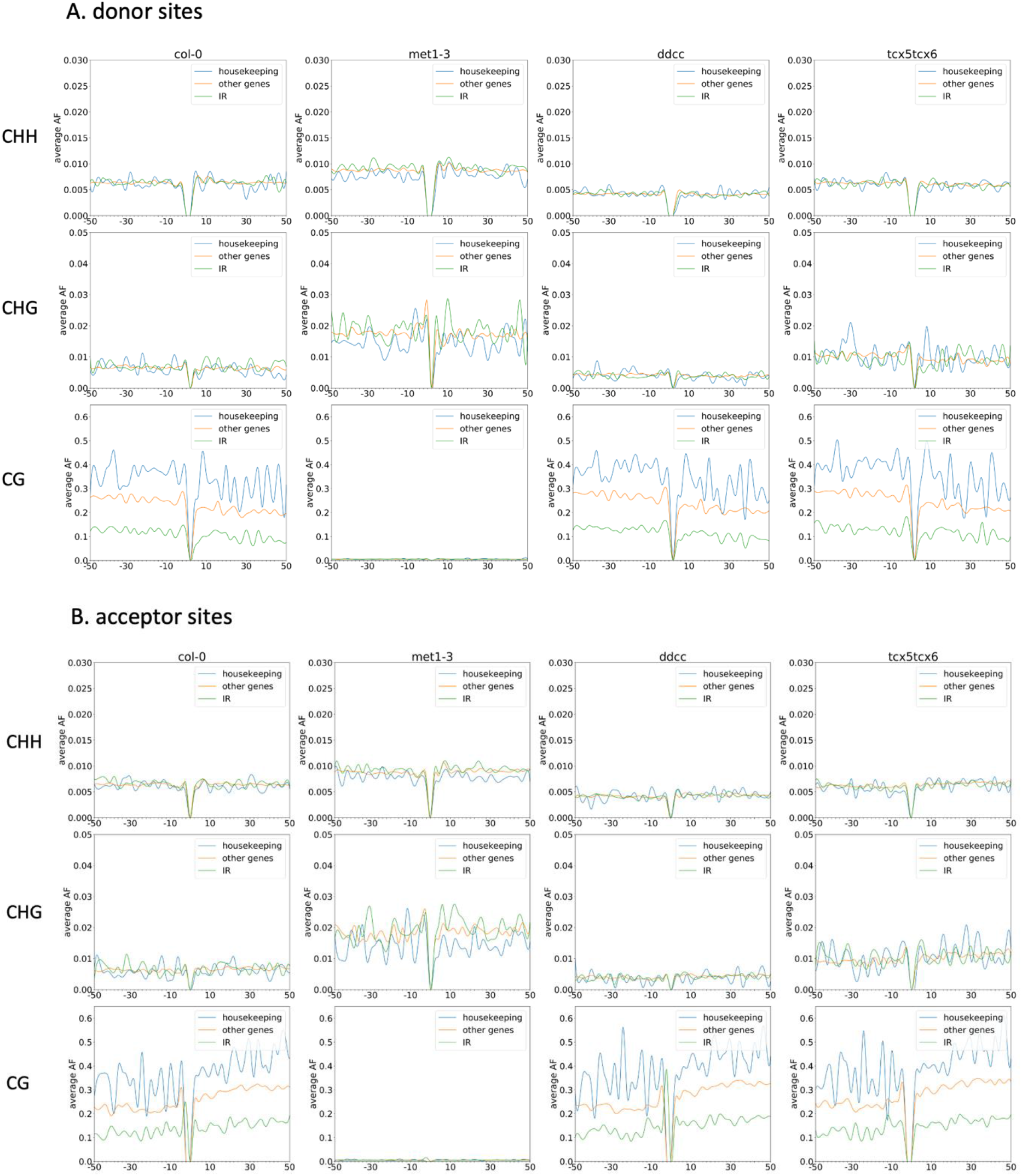
DNA methylation patterns around splicing sites in wild type (Col-0) and DNA methylation-related mutants (*met1-3, ddcc* and *tcx5tcx6*) of Arabidopsis. **A**. DNA methylation patterns around donor sites. **B**. DNA methylation patterns around acceptor sites. Splicing sites without intron retention from housekeeping genes and those from other, non-housekeeping, genes, as well as splicing sites with intron retention (IR) are shown in the CHH, CHG and CG contexts.

### ONT DRS is a powerful tool for finding novel transcripts

Using TC-RENO, we identified 117,474 transcript isoforms in all four genotypes. The number of unique transcript isoforms in each genotype is slightly different (e.g. 2,248 and 1,318 unique transcript isoforms were found in *met1-3* and *tcx5tcx6*, respectively, while only 170 and 270 unique transcript isoforms were identified in Col-0 and *ddcc*, respectively), but is far lower than the number of isoforms common to all four genotypes (77,037) (Additional file 2: Fig. S10).

The previous TC algorithm for ONT DRS reads cannot find the novel transcripts which cover a genomic region without any annotation except the chromosomal positional information. In our hands, *met1-3* activated constitutive heterochromatin are similar with such regions. To well identify those transcripts from constitutive heterochromatin, we used the TC-RENO to identified them after minimap2 (*37*) alignment, and extracted the related transcript sequence information based on *Arabidopsis* near perfect genome TAIR10 for further analysis (*38*). The advantage of ONT DRS in finding novel transcripts which may not be identified with other RNA-seq technologies, especially for transcripts that span several genes and unannotated genomic regions got embodied. To confirm these novel transcript isoforms, we chose several targets that cover several gene regions to design primers and perform nested RT-PCR to avoid false positive results.

The first target was a novel transcript containing the first exon from *At4g04810*, and the second and third exons from *At4g04830* (Fig. 5A). The second and third exon of *At4G04810*, the whole genic region of *At4G04820* and the first exon of *At4G04830* became the intron of the novel transcript. Using nested RT-PCR, we got the expected amplified fragment sizes in the first and second rounds of PCR (Fig. 5A). The second example is a novel transcript that had two exons, one from the middle exon of *At5g56010* and a second one from the last exon of *At5g56030*. The last exon of *At5g56010* and the whole genic region of *At5g56020* together with the first two exons of *At5g56030* became an intron of the novel transcript. With nested RT-PCR, the existence of this novel transcript was confirmed (Fig. 5B). We also chose a *met1-3*-specific novel transcript that had two unidentified exons distant from the other five exons which corresponded to the last five exons of *At2g22670* and confirmed it using nested RT-PCR (Fig. 5C). Another novel transcript isoform, present only in *ddcc* and *met1-3*, spanned *At2g11405, At2g11410* and *At2g11420*, but none of the exons were from these genes. Instead, all eight exons of the novel transcript were from annotated intergenic regions, with small exons and much larger introns, similar with the mammalian gene structure. This was also confirmed using nested RT-PCR (Fig. 5D). These results indicated that ONT DRS is a powerful way to find novel transcript isoforms that may be missed by other RNA-seq technologies.

**Fig. 5.**
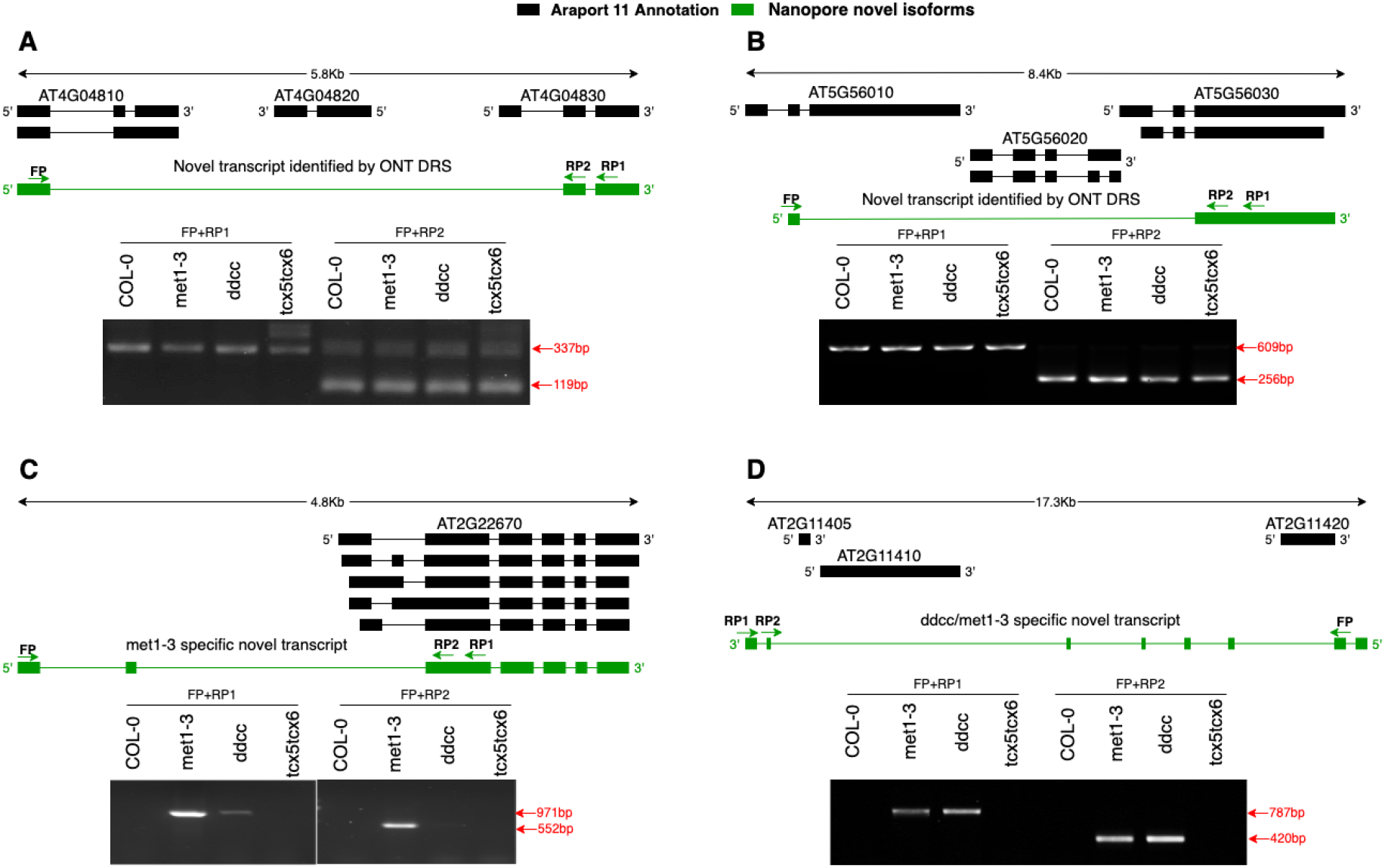
Confirmation of ONT DRS-identified novel transcript isoforms in DNA methylation-related mutants (*met1-3*/*ddcc*/*tcx5tcx6*) and wild type (Col-0) of Arabidopsis by nested RT-PCR and visualized by chemiluminescence detection. **A**. A novel transcript that combined the first exon from one gene with two final exons from another gene that was two loci downstream, with the remaining exons of these two genes and the entire genic region of the intervening gene forming an intron of this transcript. **B**. A novel transcript with the second intron of the first gene combined with the final intron of the third gene, and the intervening exons of these two genes along with the entire genic region of the intervening gene becoming the intron of this transcript. **C**. A *met1-3*-specific novel transcript containing two unidentified exons together with the five final exons of *At2g22670*. (*This gene was not labeled in panel C in the figure.) **D**. A *ddcc* and *met1-3*-specific novel transcript containing eight short exons from intergenic regions and spanning three genes. FP, forward primer; RP1, RP2, reverse primers for nested PCR. Green arrows indicate the primer orientations.

### DNA methylation may obscure potential transcription start sites (TSSs) and transcription termination sites (TTSs)

It is known that DNA methylation in some promoter regions may prevent transcription initiation, since DNA methylation-sensitive transcription factors cannot bind to their targets on the promoters to start transcription (*39*). For example, for WRKY transcription factors to bind to defense gene promoters, they need Repressor of Silencing 1 (ROS1) to demethylate DNA (*40*). Since the DNA methylation related-mutants used in this study have genome-wide effects on DNA methylation, we wanted to see if we could uncover some cryptic promoters in the genomes of these mutants by ONT DRS. Using TC-RENO, we identified a total of 89,248 TSSs among the four genotypes, with the parameter being >50 bp between two clustered TSSs, with 61,002 (>68%) of all identified TSSs being common to all four genotypes, and between 68,027 and 72,032 TSSs common to Col-0 and one of the mutants (Additional file 2: Fig. S11). The results indicated that DNA methylation had some effects on TSS. To determine these DNA methylation effects in each mutant, we analyzed the unique TSSs in each genotype. The *met1-3* mutant (with 1,472 unique TSSs) and the *tcx5tcx6* double mutant (with 1,056 unique TSSs) had more unique TSSs than the *ddcc* quadruple mutant and the wild type, with 175 and 78 unique TSSs, respectively (Additional file 2: Fig. S11). To explore whether the DNA methylation levels of the unique TSSs were lower at the promoters of the mutants than at the promoters of Col-0, we selected those unique TSSs uniquely expressed in the mutants and plotted the DNA methylation levels around these TSSs in the same regions between Col-0 and the mutants. The results showed that DNA methylation levels around these unique TSSs in *met1-3* and *ddcc* were dramatically lower than those in the same regions in Col-0 in all contexts (Fig. 6). For these TSSs in *met1-3* and *ddcc*, it is clear that DNA methylation could mask potential TSSs. However, for the *tcx5tcx6* double mutant, the difference in DNA methylation levels between Col-0 and *tcx5tcx6* was not easily discernible, although there seemed to be a very slight decrease in DNA methylation at the CHH context in *tcx5tcx6*. Whole-genome DNA methylation analysis indicated that the *tcx5tcx6* double mutant had decreased CHH methylation levels genome-wide, although CHG and CG methylation in *tcx5tcx6* was increased compared to Col-0 (Fig. 1). In addition, all the DNA methylation levels around these TSSs of *tcx5tcx6* were far lower than those around these TSSs in *met1-3* and *ddcc* (Fig. 6). These results show that the *tcx5tcx6*-specific TSSs always occurred in low DNA methylation regions, and therefore decreased CHH methylation may play a key role in transcription start. It is also possible that some specific transcription factors specifically expressed in *tcx5tcx6* are responsible for promoting transcription start in the double mutant.

**Fig. 6.**
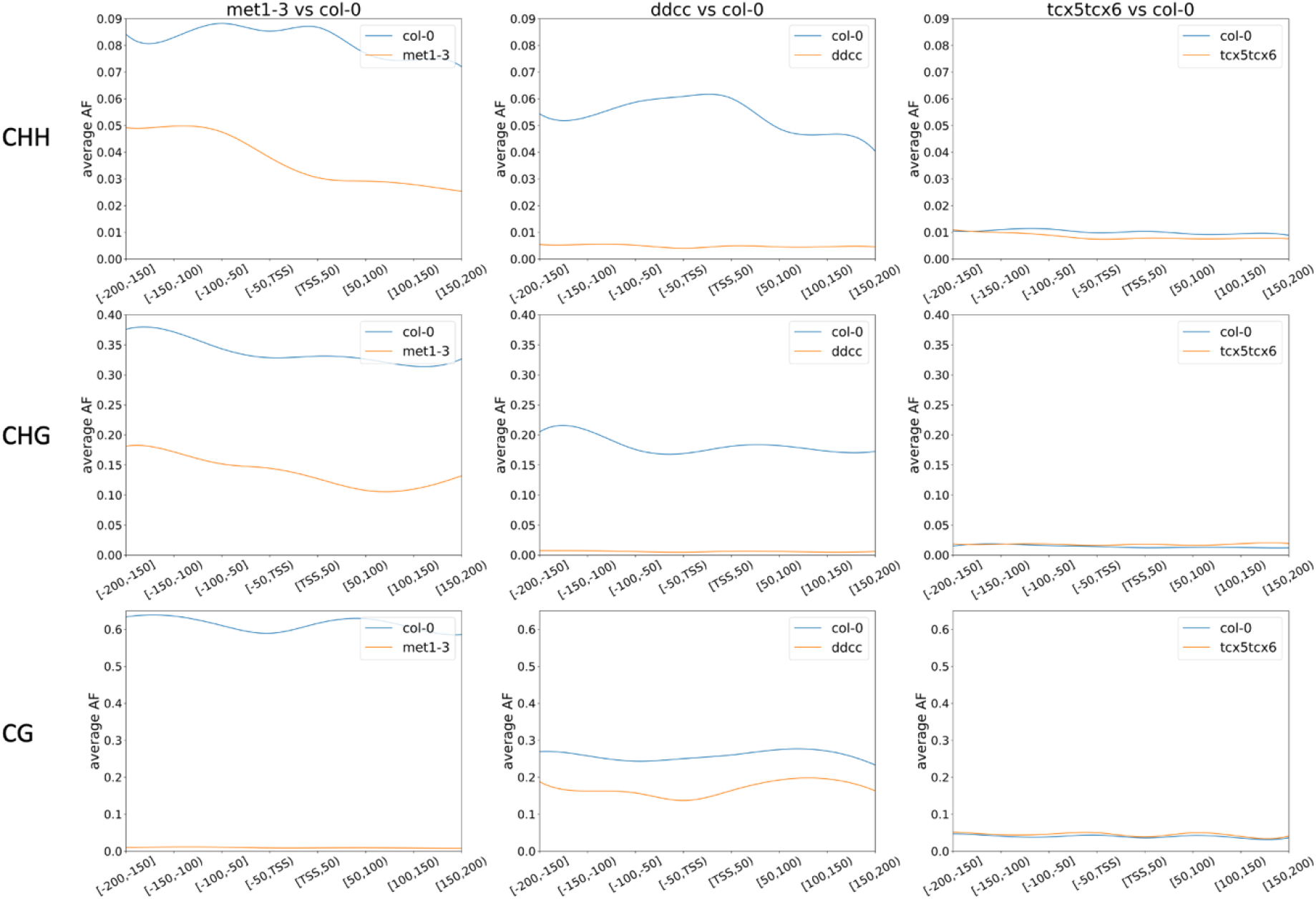
Comparisons between average DNA methylation levels in individual DNA methylation-related mutants (*met1-3*/*ddcc*/*tcx5tcx6*) and wild type (Col-0) of Arabidopsis around transcription start sites (TSSs). AF, allele frequency.

Previous studies found that altering the intronic heterochromatin status may affect the selection of alternate poly(A) sites (*41, 42*). Therefore, TTS selection could be regulated by the surrounding chromatin status that is in turn influenced by epigenetic marks including DNA methylation. It has recently been found that in mammalian cells, TTS selection is determined by DNA methylation status around the sites, since the regulator of chromatin 3-D structure, CTCF, cannot bind to methylated DNA to recruit the cohesion complex to form chromatin loops for proximal APA (alternative poly(A)) formation (*6*). Based on this information, we hypothesized that the DNA methylation level around a TTS is lower than that of its surroundings, and that a novel TTS indicates an alteration in the DNA methylation status compared to the corresponding region without a TTS. Consequently, we compared the number of TTSs among the four genotypes. TTSs common to all four genotypes (32,927) accounted for the majority of all identified TTSs (43,452), while *met1-3* had the most unique TTSs (1,472) (Additional file 2: Fig. S12). To further evaluate the correlation between DNA methylation and TTS selection, we chose transcript isoforms which bypassed a putative TTS and terminated at the TTS of the next downstream gene, being essentially fusion transcripts. By examining the DNA methylation level around the putative TTS and in the intergenic region up to the TSS of the second gene, we found that the methylation level in this stretch of DNA was significantly higher than the level in the intergenic regions of two “normal” neighboring genes with two unique transcripts (Fig. 7).

**Fig. 7.**
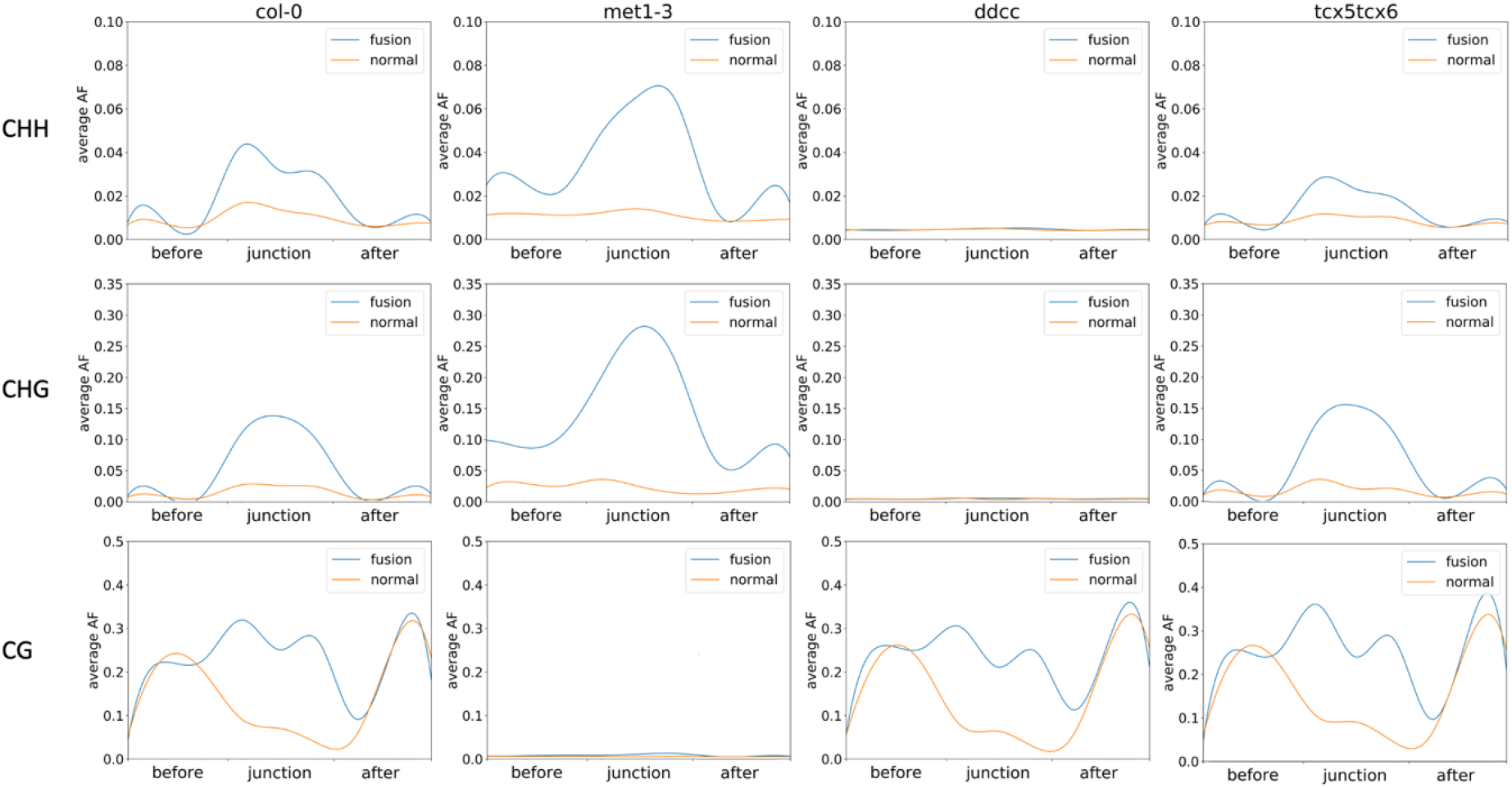
DNA methylation patterns in the intergenic regions corresponding to fusion transcripts (both cut and uncut) and normal transcripts (without readthrough) in individual DNA methylation-related mutants (*met1-3*/*ddcc*/*tcx5tcx6*) and wild type (Col-0) of Arabidopsis at the CHH, CHG and CG contexts.

### Intron-retained and fusion transcripts were detained in the nucleus

Alternative splicing is regarded as a resource of protein diversity (*43*). In mouse neurons, the profiling of ribosome-associated transcript isoforms was used to distinguish between different neurons (*44*), which shows the importance of transcript isoforms mediated by alternative splicing in development. However, another study, which profiled ribosome-associated alternatively spliced transcripts, indicated that most of the low-abundance transcripts in a transcriptome were likely not translated (*45*). Although intron retention is a frequent alternative splicing event in *A. thaliana* (*46*), the relative abundance of intron-retained transcripts is low, and only a few of them can be translated (*47*). A recent study suggested that post-transcriptionally spliced introns are not exported into the cytoplasm in plant (*7*) as detained introns reported in animal systems (*8*). Intron-retained transcripts were found to frequently introduce premature termination codons (PTCs), which may activate non-sense mediated decay (NMD) pathways (*48*). To investigate the fate of intron-retained transcripts in Arabidopsis, the subcellular distribution of several targets, including transcripts from the housekeeping gene *ACT7*, were examined to see in which cellular compartment the intron-retained transcripts were located. Our results showed that all intron-retained transcripts were detained in the nucleus irrespective of genotype (Fig. 8). The only difference among genotypes is the frequency of occurrence for the intron retention events and whether the transcripts could be further processed to produce normal transcripts. For example, there were consistently fewer reads of the intron-retained transcripts of *ACT7* and *At1g02750* in Col-0 than in other genotypes, but consistently more intron-retained transcripts of *At3g06620* in Col-0 than in *ddcc* and *tcx5tcx6* (Fig. 8). In addition, most of the intron-retained transcripts in *met1-3* were likely not further processed to form mature transcripts in the nucleus, since no intron-spliced transcripts could be found in the nucleus in *met1-3* (Fig. 8). This may be due to lower CG methylation levels in the *met1-3* mutant, and therefore not being able to slow down the transcriptional rate for mark-adding in post-transcriptional processing.

**Fig. 8.**
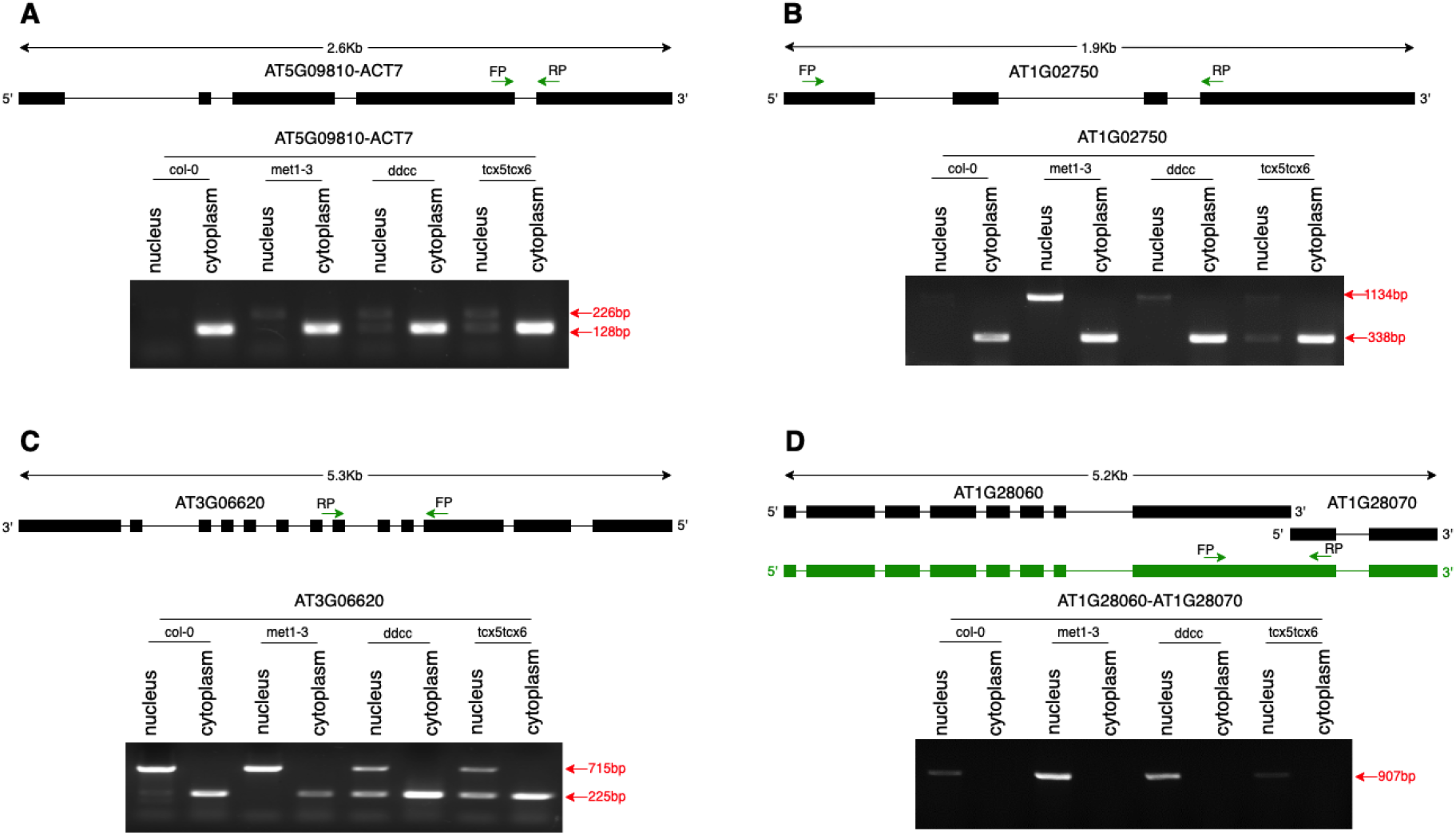
Localization of intron-retained transcripts and noncut fusion transcripts of selected genes of DNA methylation-related mutants (*met1-3*/*ddcc*/*tcx5tcx6*) and wild type (Col-0) of Arabidopsis in the nucleus and in the cytoplasm. The transcripts were first reverse-transcribed and amplified by PCR. The PCR products were separated on agarose gel and visualized by chemiluminescence detection.

Fusion transcripts of two neighboring genes is a result of the transcriptional readthrough of the upstream gene (*49*). Previously, we identified some fusion transcripts that spanned the neighboring two or more genes in Col-0 with ONT DRS (*21*). Here in the DNA methylation-related mutants, we identified more intron-retained transcripts, some of which were genotype-specific events. In addition, the intergenic regions of some of these fusion transcripts were spliced out in post-transcriptional processing while some others remained intact. Therefore, we wanted to know whether splicing out the intergenic regions is critical for exporting fusion transcripts to the cytoplasm. Therefore, we designed PCR primers to cover the intergenic regions of fusion transcripts. Our results showed that fusion transcripts with intact intergenic regions remained in the nucleus and were not exported into the cytoplasm for translation, and they were also not further processed in the nucleus (Fig. 8, Additional file 2: Fig. S13). Fusion transcripts whose intergenic regions were cut, however, were exported to the cytoplasm (Additional file 2: Fig. S13).

### DNA methylation and pre-RNA transcriptional initiation and processing in the activated constitutive heterochromatin regions in the *met1-3* mutant

Constitutive heterochromatin is highly condensed chromatin and limits the access by transcription factors (*50*), so they are normally highly silenced chromatin regions full of transposable elements and repetitive sequences (*50*). However, in *met1-3*, CG methylation was nearly completely abolished, while CHG and CHH methylation were slightly reduced (Fig. 1, Additional file 2: Fig. S2). Since DNA methylation has been linked to chromatin accessibility (*51*), its overall reduction may have enabled the constitutive heterochromatin to be activated in *met1-3* (Fig. 1, Additional file 2: Fig. S2). Although DNA methylation in the CG context was nearly eliminated in the activated constitutive heterochromatin regions, there remained a relatively high level of CHG and CHH methylation. Therefore, investigation into the relationship between DNA methylation and pre-mRNA transcriptional initiation and processing should focus specifically on the activated heterochromatin regions instead of the euchromatin regions with low DNA methylation.

Since we have already found that poly(A) tail length was positively correlated with DNA methylation, we wondered whether poly(A) tail length was also positively correlated with DNA methylation in the transcripts from constitutive heterochromatin regions. We retrieved the transcripts with poly(A) tail lengths more than 250 nt (long poly[A]) and those less than 100 nt (short poly[A]) from the activated heterochromatin regions in *met1-3* specifically and compared the DNA methylation levels around the TTS between these two groups. We found that the DNA methylation levels around the TTSs of the long poly(A) transcripts were higher than those around the TTSs of short poly(A) transcripts (Fig. 9). In addition, the poly(A) tail lengths of the transcripts from the activated heterochromatin regions in *met1-3* with very high CHG and CHH methylation were distinctively longer than those of the transcripts from the euchromatin regions that could also be transcribed in Col-0 (Additional file 2: Fig. S14). The results confirmed the positive correlation between DNA methylation and poly(A) tail length.

**Fig. 9.**
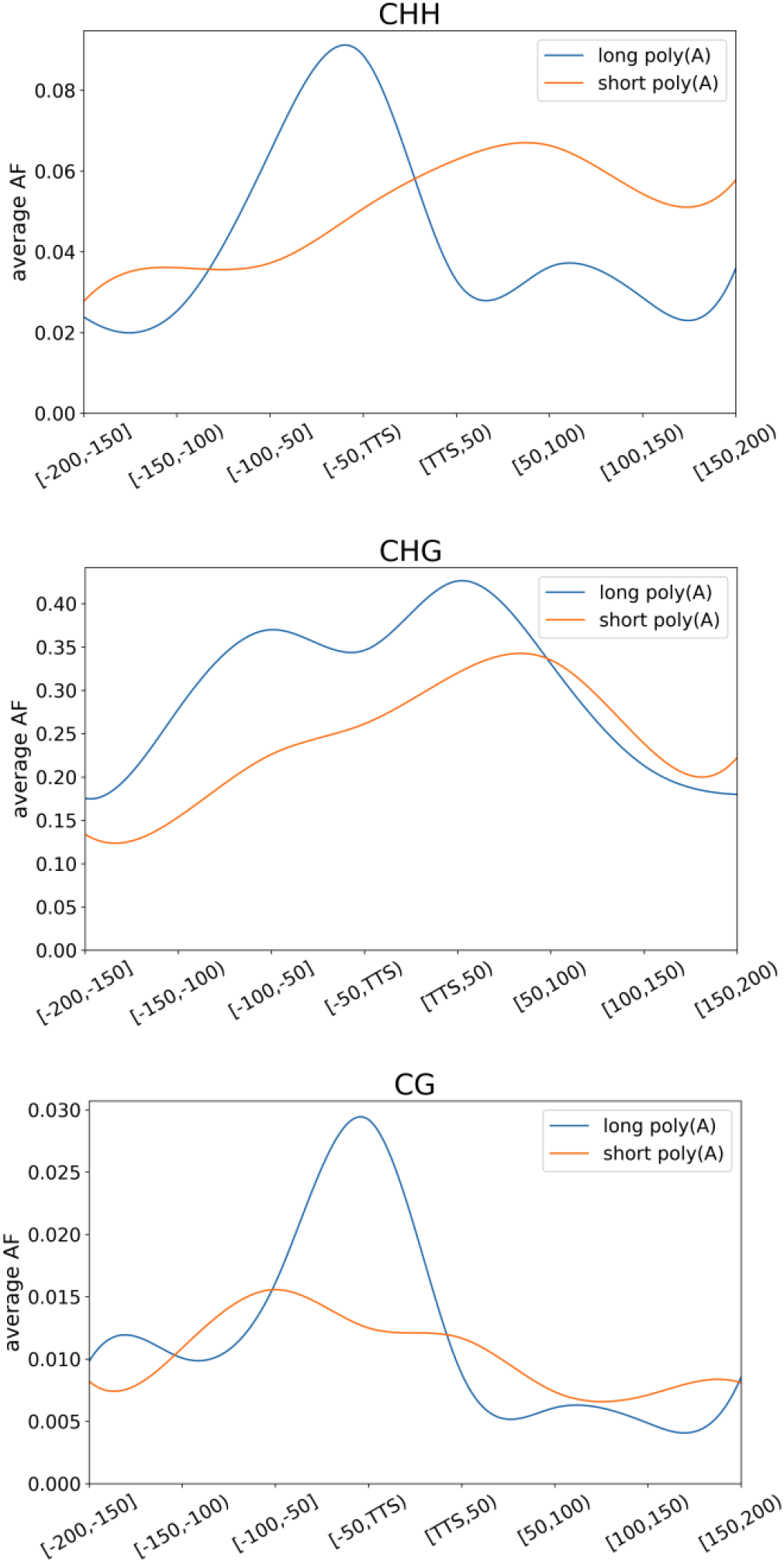
DNA methylation patterns around transcription termination sites (TTSs) of transcripts with a poly(A) tail longer than 250nt (long poly[A]) and with a poly(A) tail length shorter than 100nt (short poly[A]) from activated constitutive heterochromatin of the DNA methylation-deficient mutant, *met1-3*, of Arabidopsis at the CHH, CHG and CG contexts. AF, allele frequency.

Since the DNA methylation levels around the splicing sites of intron-retained transcripts were found to be lower than those around fully spliced sites earlier, we then wanted to know whether the splicing sites (both donor and acceptor sites) of the intron-containing transcripts from the activated constitutive heterochromatin regions of *met1-3* also had lower DNA methylation levels. The results showed that DNA methylation around splicing sites of intron-containing transcripts did have lower CHG and CHH methylation levels than around fully spliced sites in the corresponding transcripts (Fig. 10). The results confirmed that lower DNA methylation around splicing sites may be the cause of intron retention events.

**Fig. 10.**
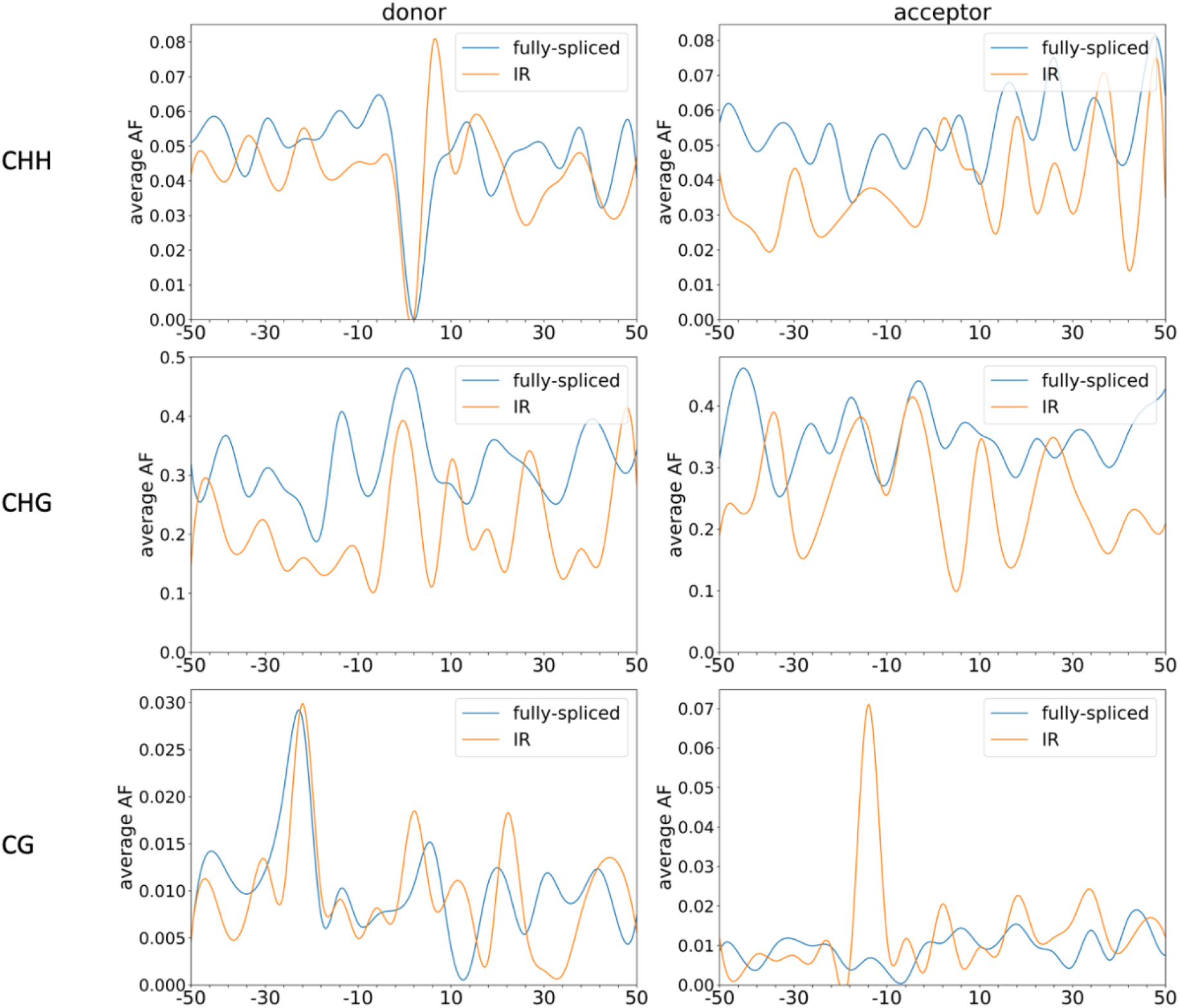
DNA methylation levels around the splicing sites of fully-spliced versus intron-retained (IR) transcripts from the activated constitutive heterochromatic regions of the DNA methylation-deficient mutant, *met1-3*, of Arabidopsis at the CHH, CHG and CG contexts.

## Discussion

### DNA methylation and poly(A) tail length

The poly(A) tail may facilitate mature mRNA export from the nucleus to the cytoplasm (*32*) and promote translational efficiency (*52*). Interestingly, our results showed that the transcripts from TE genes had much longer poly(A) tails than those from other genes. Normally, poly(A) is added immediately after the TTS during 3’ end processing that takes place concurrently with transcription (*13*), while the chromosomal regions encoding TEs are always hypermethylated. A previous study showed that poly(A) polymerase 1 (PAPS1) can interact with the RNA-directed DNA methylation (RdDM) pathway (*53*) which mediates *de novo* CHH methylation. Our results showed that CHH methylation levels were highest around the TTSs of TE genes and lowest around those of housekeeping genes, especially in the *met1-3* mutant (Fig. 3). Therefore, some RdDM pathway components may sustain PAPS1 to complete co-transcriptional processing, thus facilitating poly(A) tail formation. Alternatively, DNA methylation around TTSs may prevent PAPS1 from escaping from the chromatin scaffold. However, poly(A) tails can also be added by the non-canonical poly(A) polymerases, TRF4 and TRF5, in yeast (*54*), with the resulting RNAs subjected to degradation via exosome (*54*). *A. thaliana* has a *TRF4/5* homolog, *TRL* (*At5g53770*), which has been suggested to be able to add a poly(A) tail for a variety of RNAs for degradation via exosome (*55*). Therefore further investigation is required to determine whether TE transcripts with a longer poly(A) tail tend to be degraded.

### DNA methylation, intron retention, and alternative splicing site selection

Alternative splicing is a common mechanism for regulating gene expression (*56*), with intron retention being one of the most common type (*36*). The retained introns in mRNAs have been further classified as either non-sense-mediated decay pathway (NMD)-sensitive or NMD-resistant. For example, retained introns in mRNAs in animal cells (*8*) and post-transcriptionally spliced introns in plant mRNAs (*7*) are resistant to NMD. NMD-resistant introns were not detected in the cytoplasm in this study, since they were retained in the nucleus as was confirmed with RT-PCR (Fig. 8). Since pre-mRNA processing, including alternative splicing, is co-transcriptionally regulated, chromatin accessibility may affect the transcription rate, as well as co-transcriptional splicing (*2*). In *A. thaliana*, although the DNA methylation levels in exons are considerably lower than those in mammalian exons (*57*), CG methylation in the gene body is also generally higher than in the surrounding chromatin, especially for housekeeping genes (*58*). In plants, constitutively transcribed genes normally have hypermethylated exons, and they are generally not inducible or development-related genes (*58*). Here we found that around the putative splicing sites, CG methylation was always lower at the intron-retained transcripts than at the fully spliced transcripts of housekeeping and other genes (Fig. 4), which supports a previous hypothesis that DNA methylation around splicing sites may slow down transcription (*4*), and thus give spliceosome enough time to find the splicing sites to splice the introns. These results indicated that DNA methylation is important for splicing site selection, either via slowing down the transcriptional rate with a roadblock created by a methylated DNA binding protein, such as meCP2 (*5*), to facilitate splicing site targeting by the spliceosome, or via a spliceosome-interacting heterochromatin protein, such as the mammalian HP1, to recruit splicing factors (*59*).

### Intron retention and the export of mRNA from nucleus to cytoplasm

It is known that transcripts with retained introns or post-transcriptionally spliced introns cannot be exported to the cytoplasm (*7, 8*), but some intron-containing transcripts can be processed post-transcriptionally to remove the intron and therefore are able to be exported from the nucleus (*8*). However, here we observed that some intron-containing transcripts had a lower chance of being further processed in the nucleus to remove the intron post-transcription, especially in *met1-3* where CG methylation was almost completely eliminated (Fig. 8). One possible explanation for this variation is that low or no DNA methylation in the *met1-3* mutant may result in quicker transcription, leaving no time to form a junction complex to mark the splicing sites for further processing by a spliceosome after transcription, and the intron is therefore unspliced. On the other hand, in the *tcx5tcx6* mutant with increased CHG and CG methylation, intron-containing transcripts could more easily be further processed in the nucleus (Fig. 8). Fusion transcripts with an unspliced intergenic region may be regarded as intron-containing transcripts, and thus cannot be exported to the cytoplasm. Since the intergenic region does not have putative splicing sites, it cannot be marked by factors to be recognized by the post-transcriptional splicing complex. Therefore, these fusion transcripts cannot be further processed in the nucleus in order to be exported to the cytoplasm (Fig. 8D, Additional file 2: Fig. S13). At the same time, fusion transcripts with spliced-out intergenic regions were regarded as normal transcripts and thus could be exported from the nucleus (Additional file 2: Fig. S13).

### ONT DRS can detect novel transcripts, especially those from highly silenced genomic regions

ONT DRS can sequence native RNA molecules without being limited by the length of the sequenced RNA molecules. Here, RNA molecules longer than 13,000 nt transcribed from the locus *At4g17140* were sequenced, and, using the ONT DRS results, the misannotated splicing sites were corrected (*21*). Using ONT DRS, we could obtain full-length, corrected transcript isoforms in Col-0 and *met1-3*. Hence ONT DRS is a powerful tool for finding very long full-length RNA molecules which may not be detected using other sequencing technology. Other RNA-seq technologies need to covert RNA molecules to cDNA, where the maximal reverse-transcribable length of RNA molecules by current, commercially available, reverse transcriptases is less than 12,000 nt. ONT DRS is also a powerful technology for finding hidden transcripts that have never been reported because their DNA templates cover large genomic regions, and their gene structures do not resemble the typical plant gene structure. An example is the novel transcript present in *met1-3* and *ddcc*, which spanned *At2g11405, At2g11410* and *At2g11420*, as well as the large intergenic region of *At2g11410* and *At2g11420* (Fig. 5). The gene structure of this novel transcript resembles the mammalian gene structure, with very long introns and very short exons (Fig. 5).

Constitutive heterochromatin regions, which consist of TEs and repetitive sequences, are tightly silenced in the wild type. However, in *met1-3*, the repressive epigenetic mark of CG methylation is nearly eliminated and therefore the chromatin status of these regions is altered (*51*), thus giving access to transcription factors and DNA-dependent RNA polymerase II to initiate transcription. As it was pointed out previously, short-read sequencing technologies would not be able to recognize transcripts of TE genes or those with multiple repetitive sequences. On the other hand, ONT DRS can sequence full-length native RNA molecules regardless of transcript lengths or sequences, so it provides a good solution for detecting the transcript isoforms from these heterochromatin regions, and opens a window into the relationship between DNA methylation and pre-mRNA transcriptional initiation and processing. Although observations on the transcript isoforms from the activated constitutive heterochromatin regions are complicated by the presence of CHG and CHH hypermethylation in *met1-3*, the conclusions we drew with respect to transcriptional initiation, intron retention and poly(A) tail length of transcripts from the activated constitutive heterochromatin regions of *met1-3* are still consistent with those we made with respect to the euchromatin regions.

## Conclusions

This study elucidated the relationship between DNA methylation and pre-mRNA transcriptional initiation and processing. Transcription initiation requires hypomethylation of the promoter regions, while lower DNA methylation around splicing sites tend to cause intron retention events. DNA methylation around the TTS correlates well with longer poly(A) tail length. All these observations regarding the relationship between DNA methylation and pre-mRNA transcriptional initiation and processing were made possible with ONT DRS in combination with BS-seq, and further confirmed with transcripts from the activated constitutive heterochromatin regions in the DNA methylation-deficient mutant, *met1-3*.

## Methods

### Plant materials

Seeds of *Arabidopsis thaliana* wild type Col-0 seeds and DNA methylation-related mutants, *met1-3, ddcc* (*3*) and *tcx5tcx6* (*25*), were surface-sterilized with 50% commercial bleach and 0.01% Triton X-100 for 10 minutes, and then washed 5 times with sterilized H_2_O. The sterilized seeds were sown on ½ Murashige and Skoog (MS) plates with 1% sucrose and 0.6% agar (Millipore-Sigma cat# A1296). After stratification at 4°C for 48 hours, they were placed in a growth chamber with 16 h light/8 h dark for 12 days. The seeds of *met1-3* (Arabidopsis Biological Research Center - ABRC) are heterozygous, from which we generated first-generation homozygous seeds for this study. The seeds for the *ddcc* quadruple mutant were provided by Professor Xiaofeng Cao (Institute of Genentics and Developmental Biology, Chinese Academy of Sciences), and those for the *tcx5tcx6* double mutant were a gift from Dr. Xinjian He (National Institute of Biological Sciences, Beijing).

### DNA/RNA extraction

Twelve-day-old seedlings of col-0, *met1-3, ddcc* and *tcx5tcx6* were harvested and ground to a powder in liquid nitrogen, and genomic DNA were extracted with GeneJET Plant Genomic DNA purification kit (ThermoFisher, Cat# K0791) and RNA were extracted with Trizol.

For total RNA extraction, 1 ml Trizol was added to each 100 mg of frozen ground plant material and mixed well. The solution was incubated at room temperature for 5 min, and then 200 μL chloroform was added. The mixture was incubated at room temperature for an additional 3 min, before being centrifuged at 4°C at 12,000 *g* for 10 min, after which the supernatant was transferred to a fresh tube and mixed with an equal volume of isopropanol at room temperature on the HulaMixer (Thermo Fisher Scientific) at low speed for 10 min. The RNA was pelleted via centrifugation at 4°C at 12,000 *g* for 15 min. After washing with 75% ethanol, 100 μL RNase-free H_2_O was used to dissolve the RNA pellet. The resulting RNA was digested with RNase-free DNase I (NEB, M0303S), and purified with acidic phenol (125:24:1 (phenol:chloroform:isoamyl), pH4.5, Thermo Fisher Scientific, Cat# AM9722) via Phase Lock Gel light (5PRIME, Cat# 2302820). The aqueous supernatant was transferred to a fresh tube and mixed with lithium chloride to a final concentration of 2.5 M. The mixture was incubated at -20 °C for 1 h, and then it was centrifuged at 4 °C for 30 min at 12,000 *g*.

The resulting RNA pellet was washed twice with 75% ethanol, and then dissolved in RNase-free H_2_O. For poly(A) RNA extraction, 75 μg of purified total RNA in 100 μL RNase-free H_2_O was used to isolate mRNAs with Dynabeads™ mRNA Purification Kit (Thermo Fisher Scientific, Cat# 61006) following the manufacturer’s manual.

### RNA extraction from the nuclear and cytoplasmic fractions of the same sample

Total RNA extraction from the nuclear and cytoplasmic fractions of the same plant sample was carried out according to a published protocol (*60*). Briefly, 1 g of tissue from X-day-old Arabidopsis seedlings was ground into a powder in liquid nitrogen, and then mixed with 2 mL lysis buffer (20 mM Tris-HCl, pH7.5, 20 mM KCl, 2 mM EDTA, 2.5 mM MgCl_2_, 25% glycerol, 250 mM sucrose, 5 mM DTT and 1/100X protease inhibitor cocktail, [Millipore-Sigma, Cat# P9599]). After filtration with 2 layers of Miracloth, the flowthrough was centrifuged at 1,500 *g* at 4°C for 10 min, and the supernatant was transferred to a fresh tube and further centrifuged at 10,000 *g* for 10 min at 4°C. The resulting supernatant was collected as the cytoplasmic fraction and was extracted with Trizol to isolate cytoplasmic RNAs for RT-PCR. The pellet from the initial 1,500 *g* spin was washed 8 times with 8 mL each nuclear suspension buffer 1 (20 mM Tris-HCl, pH7.4, 2.5 mM MgCl_2_ and 0.2% Triton X-100), and then resuspended with 500 μL nuclear resuspension buffer 2 (20 mM Tris-HCl, pH7.5, 10 mM MgCl_2_, 250 mM sucrose, 0.5% Triton X-100, 5 mM 2-mercaptoethanol and 1X protease inhibitor cocktail [Millipore-Sigma, Cat# P9599]), and carefully overlaid on top of 500 μL of nuclear resuspension buffer 3 (20 mM Tris-HCl, pH7.5, 10 mM MgCl_2_, 1.7 M sucrose, 0.5% Triton X-100, 5 mM 2-mercaptoethanol and 1x protease inhibitor cocktail [Millipore-Sigma, Cat# P9599]). The sample was centrifuged at 16,000 *g* at 4°C for 45 min. The final pellet was used for nuclear RNA extraction with Trizol.

### ONT DRS library construction and sequencing

900ng purified mRNAs from each sample were used to construct an ONT DRS library with Nanopore ONT DRS SQK-RNA002 kit. The manual for ONT DRS library construction was strictly followed except that the RNA CS was replaced with mRNA from samples. The constructed ONT DRS library was loaded onto FLO-MIN106 flowcell (R9.4.1), and run in the MinION sequencing platform.

### Nested RT-PCR for novel transcript and transcript variant identification

For each sample, 1 μg of total RNA was used to perform reverse transcription (RT) with *EasyScript*® First-Strand cDNA Synthesis SuperMix (TransGen Biotech, Cat# AE301-02) for novel transcript identification. The resulting cDNA was used for two rounds of nested PCR. The first round was performed with forward primer and reverse primer 1 (RP1), following protocol X (*include the volumes of each component: primers, cDNA and PCR reagents, plus the PCR steps). Then 1 μL of the PCR product was diluted with 30 μL distilled deionized H_2_O (dd H_2_O). Then 1 μL of the diluted mixture was used as a template for the second round of PCR with forward primer and RP2, following this protocol. Then the first and second-round PCR products were resolved in a 1.7% agarose gel and stained with Gel red dye and documented with the Biorad Chemidoc system.

For RT-PCR of nuclear RNAs and cytoplasmic RNAs, an equal amount of total RNA from the cytoplasmic fraction of each genotype, and an equal amount of total RNA from the nuclear fraction of each genotype, was used. The amount of cytoplasmic RNA used was two times that of the nuclear RNA. The RT and PCR reactions were performed as described above.

### Bisulfite sequencing (BS-seq)

The BS-seq library was constructed by BGI Company. Briefly, 1 μg of genomic DNA from each sample (col-0, *ddcc* and *met1-3*) was fragmented to 200-300 bp, and the fragmented genomic DNA was confirmed in 1.5% agarose gel, and purified with the MiniElute PCR purification kit (QIAGEN, Cat. No. / ID: 28006×4). The purified genomic DNA fragments were end-repaired and had an A-tail added to the 3’ end. Then the A-tailed DNA fragments were ligated with a methylated adapter sequence and purified again with the QIAGEN MiniElute PCR purification kit. The purified adapter-added genomic DNA was subjected to bisulfite conversion using the EZ DNA Methylation-Gold kit (Zymo Research, Cat#: D5005). Then the treated genomic DNA was resolved in a 1.5% agarose gel, and fragments in the 320 to 420 bp range were purified with the QIAquick Gel extraction kit (QIAGEN, Cat#:28706). The purified genomic DNA fragments were amplified by several rounds of PCR, and the resulting PCR products of 320-420 bp in length were resolved in a 1.5% agarose gel and purified for the BS-seq library. After validation of the library, they were sequenced on the BGI sequencing platform.

### Base-calling and pre-filtering of Nanopore long reads

The raw reads from ONT DRS were base-called using Guppy basecaller (version 3.4.4) (*61*) with the high accuracy model (rna_r9.4.1_70bps_hac.cfg). Low-quality reads with the Phred quality score below 7 were removed. The *polya* subprogram of nanopolish (version 0.13.2) (*62*) was used to filter out reads with truncated 3′ ends (reads with QC tags “READ_FAILED_LOAD”, “SUFFCLIP”, or “NOREGION”), and to estimate the poly(A) tail length. The filtered reads were randomly sub-sampled to a similar depth of coverage before all the reads were combined for transcriptome reconstruction. The reads were mapped to the Araport11 reference genome using Minimap2 (version 2.17-r941) (*37*) with the parameters “-ax splice -k14 -uf --junc-bed --junc-bonus 9”. Only unique alignments were considered for further analyses.

### Transcript isoform identification and quantification analysis

In order to adapt to the purpose of our study, the tool TC-RENO was optimized using the basics of TrackCluster (*63*) with modifications in the clustering method and the addition of novel functions. The detailed TC-RENO designs and the associated codes are available on GitHub (https://github.com/HKU-BAL/TC-RENO).

The isoforms were detected by TC-RENO with default parameters. Their supported read counts provided in the output were then used to compute the CPM (counts-per-million) per isoform in each sample using edgeR (version 3.24.2) (*64*).

For visualization, the per-site expression level was calculated by summing the CPM of its supported isoforms. It was then used to obtain an average within each non-overlapping 20,000-bp sliding window based on Araport11. After pairwise comparison between each mutant and Col-0, the differential expression regions (DERs), represented as log_2_ fold-change and defined as the difference between the average CPM per window, were presented in a chromosome-level plot.

### Genome-wide methylation-calling using bisulfite sequencing data

The paired-end whole-genome bisulfite sequencing data were aligned to the Araport11 reference genome and the methylated 5mCs (5-Methylcytosine) were extracted using the default setting “-p -- no_overlap --comprehensive --cytosine_report” of Bismark (version 0.22.3) (*65*). For each sample, the CG, CHG and CHH Bismark outputs were formatted into separate BigWig files, including methylated genomic positions, methylation allele frequencies (MAF) and strand information for downstream analyses.

Similar to the calculation of DERs, the differential methylation regions (DMRs) were calculated using the corresponding average MAF for CG, CHG and CHH per non-overlapping 20,000-bp sliding window in each sample, and were presented in a chromosome-level plot.

### Discovery of alternative splicing events

The regions within 25 bp, 50 bp, 5 bp and 5 bp of the transcription start sites (TSSs), transcription termination sites (TTSs), and acceptor and donor sites of mRNA isoforms, respectively, were separately merged into clusters. To determine intron retention (IR), exon skip and extra exon events, novel isoforms were mapped to genes in the reference genome. Isoforms with ≥ 50% of exons intersecting with the exons of the gene in Araport11 were regarded as a member of the gene. If an isoform intersected with multiple genes, the similarity score was computed as:

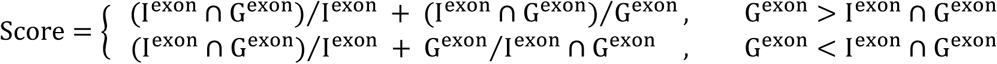

between the isoform, I, and the gene, G, such that the isoform was assigned to the gene with the highest similarity score. Subsequently, the exons and introns of each novel isoform were compared against the TAIR-defined canonical transcript of the assigned genes to detect alternative splicing events.

### Detection of *met1-3* activated constitutive heterochromatin regions

The *met1-3* activated constitutive heterochromatin regions were defined as the transcribed regions in the *met1-3* genome with high CG MAFs that were not found in Col-0. The isoforms expressed in *met1-3* that did not intersect with the reads from Col-0 were extracted, except for the isoforms with ultra-long introns (>10,000 bp). The remaining isoforms were intersected with each other using bedtools (version 2.29.2 (*66*) with the parameters “-s -f 0.5”. The overlapping isoforms were clustered together to form a transcribed region. The first TSS and the last TTS were defined as the start and end positions of the region respectively. Regions with >0.8 of the average CG MAF of Col-0 on the coding strand were considered to be *met1-3* activated constitutive heterochromatin regions.

### Poly(A) tail length analysis

To compare the poly(A) tail length of reads at the gene level, the same method for isoform assignment as described above was applied to assign reads to genes. Then, to compare the poly(A) length between the heterochromatin and euchromatin regions of *met1-3*, the *met1-3* reads were identified in the activated constitutive heterochromatin regions if at least 50% of their alignment overlapped with the regions. Furthermore, the TTS sites of the reads in the *met1-3* activated consitutive heterochromatin regions were sorted according to whether their poly(A) tails were longer than 250 nt (i.e. long poly[A]) or shorter than 100 nt (i.e. short poly[A]). The TTS sites shared by both long and short poly(A) reads, or sites with no more than 3 supporting reads were excluded. The MAF distribution patterns on the coding strand were compared between the two types of TTS sites.

### Distribution of MAFs in fusion and normal genes

A transcript was annotated as a fusion if it overlapped with at least 50% of one independent exon in two or more genes on the same strand of the same chromosome. The genes associated with the identified fusion transcripts were defined as fusion genes, while two adjacent expressed genes with a gap of fewer than 1,000 bp and not covered by fusion transcripts were considered to be normal genes. The MAF on the coding strand between the normal and fusion genes was compared.

## Declarations

### Availability of data

Our data is available in gene expression omnibus (GEO) at GSE172010.

### Authors’ contributions

S.Z., R.L. and H.-M. L. conceived the project, designed the overall research strategy, and coordinated the data collection and analyses. S.Z., S.C., Y.L. and L.Z. conducted the experiments. R.L., Q.L., W.S.L. and Y.X. performed the data analyses and bioinformatics. S.Z., R.L., Q.L., W.S.L. and H.-M. L. co-wrote the first draft of the manuscript. All authors were involved in the preparation of the final manuscript.

## Acknowledgements

We thank Dr. Xiaofeng Cao (Institute of Genetics and Developmental Biology, Chinese Academy of Sciences) and Dr. Xinjian He (National Institute of Biological Sciences, Beijing) for sharing with us the ddcc quadruple mutant and tcx5tcx6 double mutant used in this study. We also thank Cathy Man, Iris Tong and Man Wah Li at CUHK for their help with administration and facility access and Ms. Jee Yan Chu copy-edited this manuscript.

## References

1. D. L. Bentley, Coupling mRNA processing with transcription in time and space. Nat Rev Genet 15, 163–175 (2014).

2. S. J. Brown, P. Stoilov, Y. Xing, Chromatin and epigenetic regulation of pre-mRNA processing. Hum Mol Genet 21, R90–96 (2012).

3. Q. Qiu et al., DNA methylation repels targeting of Arabidopsis REF6. Nat Commun 10, 2063 (2019).

4. S. Shukla et al., CTCF-promoted RNA polymerase II pausing links DNA methylation to splicing. Nature 479, 74–79 (2011).

5. J. I. Young et al., Regulation of RNA splicing by the methylation-dependent transcriptional repressor methyl-CpG binding protein 2. Proc Natl Acad Sci U S A 102, 17551–17558 (2005).

6. V. Nanavaty et al., DNA Methylation Regulates Alternative Polyadenylation via CTCF and the Cohesin Complex. Mol Cell 78, 752–764 e756 (2020).

7. J. Jia et al., Post-transcriptional splicing of nascent RNA contributes to widespread intron retention in plants. Nat Plants 6, 780–788 (2020).

8. P. L. Boutz, A. Bhutkar, P. A. Sharp, Detained introns are a novel, widespread class of post-transcriptionally spliced introns. Genes Dev 29, 63–80 (2015).

9. M. Dreyfus, P. Regnier, The poly(A) tail of mRNAs: bodyguard in eukaryotes, scavenger in bacteria. Cell 111, 611–613 (2002).

10. S. A. Lima et al., Short poly(A) tails are a conserved feature of highly expressed genes. Nat Struct Mol Biol 24, 1057–1063 (2017).

11. A. O. Subtelny, S. W. Eichhorn, G. R. Chen, H. Sive, D. P. Bartel, Poly(A)-tail profiling reveals an embryonic switch in translational control. Nature 508, 66–71 (2014).

12. S. M. Bresson, N. K. Conrad, The human nuclear poly(a)-binding protein promotes RNA hyperadenylation and decay. PLoS Genet 9, e1003893 (2013).

13. U. Kuhn et al., Poly(A) tail length is controlled by the nuclear poly(A)-binding protein regulating the interaction between poly(A) polymerase and the cleavage and polyadenylation specificity factor. J Biol Chem 284, 22803–22814 (2009).

14. W. M. Freeman, S. J. Walker, K. E. Vrana, Quantitative RT-PCR: pitfalls and potential. Biotechniques 26, 112-122, 124-115 (1999).

15. G. Zhang et al., PacBio full-length cDNA sequencing integrated with RNA-seq reads drastically improves the discovery of splicing transcripts in rice. Plant J 97, 296–305 (2019).

16. V. Potapov et al., Base modifications affecting RNA polymerase and reverse transcriptase fidelity. Nucleic Acids Res 46, 5753–5763 (2018).

17. C. K. Kwok, G. Marsico, A. B. Sahakyan, V. S. Chambers, S. Balasubramanian, rG4-seq reveals widespread formation of G-quadruplex structures in the human transcriptome. Nat Methods 13, 841–844 (2016).

18. Y. Ding et al., In vivo genome-wide profiling of RNA secondary structure reveals novel regulatory features. Nature 505, 696–700 (2014).

19. D. R. Garalde et al., Highly parallel direct RNA sequencing on an array of nanopores. Nat Methods 15, 201–206 (2018).

20. C. Soneson et al., A comprehensive examination of Nanopore native RNA sequencing for characterization of complex transcriptomes. Nat Commun 10, 3359 (2019).

21. S. Zhang et al., New insights into Arabidopsis transcriptome complexity revealed by direct sequencing of native RNAs. Nucleic Acids Res 48, 7700–7711 (2020).

22. M. T. Parker et al., Nanopore direct RNA sequencing maps the complexity of Arabidopsis mRNA processing and m(6)A modification. Elife 9, (2020).

23. G. Almouzni, H. Cedar, Maintenance of Epigenetic Information. Cold Spring Harb Perspect Biol 8, (2016).

24. S. W. Chan, I. R. Henderson, S. E. Jacobsen, Gardening the genome: DNA methylation in Arabidopsis thaliana. Nat Rev Genet 6, 351–360 (2005).

25. Y. Q. Ning et al., DREAM complex suppresses DNA methylation maintenance genes and precludes DNA hypermethylation. Nat Plants 6, 942–956 (2020).

26. H. Stroud, M. V. Greenberg, S. Feng, Y. V. Bernatavichute, S. E. Jacobsen, Comprehensive analysis of silencing mutants reveals complex regulation of the Arabidopsis methylome. Cell 152, 352–364 (2013).

27. L. Sun et al., TDNAscan: A Software to Identify Complete and Truncated T-DNA Insertions. Front Genet 10, 685 (2019).

28. H. Wollmann et al., The histone H3 variant H3.3 regulates gene body DNA methylation in Arabidopsis thaliana. Genome Biol 18, 94 (2017).

29. A. Teissandier, D. Bourc’his, Gene body DNA methylation conspires with H3K36me3 to preclude aberrant transcription. EMBO J 36, 1471–1473 (2017).

30. B. J. Natalizio, S. R. Wente, Postage for the messenger: designating routes for nuclear mRNA export. Trends Cell Biol 23, 365–373 (2013).

31. F. F. Aceituno, N. Moseyko, S. Y. Rhee, R. A. Gutierrez, The rules of gene expression in plants: organ identity and gene body methylation are key factors for regulation of gene expression in Arabidopsis thaliana. BMC Genomics 9, 438 (2008).

32. L. A. Castellano, A. A. Bazzini, Poly(A) tails: longer is not always better. Nat Struct Mol Biol 24, 1010–1011 (2017).

33. Z. Dominski, X. C. Yang, W. F. Marzluff, The polyadenylation factor CPSF-73 is involved in histone-pre-mRNA processing. Cell 123, 37–48 (2005).

34. C. R. Mandel et al., Polyadenylation factor CPSF-73 is the pre-mRNA 3’-end-processing endonuclease. Nature 444, 953–956 (2006).

35. E. Wahle, W. Keller, The biochemistry of 3’-end cleavage and polyadenylation of messenger RNA precursors. Annu Rev Biochem 61, 419–440 (1992).

36. H. Ner-Gaon et al., Intron retention is a major phenomenon in alternative splicing in Arabidopsis. Plant J 39, 877–885 (2004).

37. H. Li, Minimap2: pairwise alignment for nucleotide sequences. Bioinformatics 34, 3094–3100 (2018).

38. C. Y. Cheng et al., Araport11: a complete reannotation of the Arabidopsis thaliana reference genome. Plant J 89, 789–804 (2017).

39. Y. A. Medvedeva et al., Effects of cytosine methylation on transcription factor binding sites. BMC Genomics 15, 119 (2014).

40. T. Halter et al., The Arabidopsis active demethylase ROS1 cis-regulates defence genes by erasing DNA methylation at promoter-regulatory regions. Elife 10, (2021).

41. M. Lei et al., Arabidopsis EDM2 promotes IBM1 distal polyadenylation and regulates genome DNA methylation patterns. Proc Natl Acad Sci U S A 111, 527–532 (2014).

42. L. Ma, C. Guo, Q. Q. Li, Role of alternative polyadenylation in epigenetic silencing and antisilencing. Proc Natl Acad Sci U S A 111, 9–10 (2014).

43. D. L. Black, Protein diversity from alternative splicing: a challenge for bioinformatics and post-genome biology. Cell 103, 367–370 (2000).

44. E. Furlanis, L. Traunmuller, G. Fucile, P. Scheiffele, Landscape of ribosome-engaged transcript isoforms reveals extensive neuronal-cell-class-specific alternative splicing programs. Nat Neurosci 22, 1709–1717 (2019).

45. R. J. Weatheritt, T. Sterne-Weiler, B. J. Blencowe, The ribosome-engaged landscape of alternative splicing. Nat Struct Mol Biol 23, 1117–1123 (2016).

46. Y. Marquez, J. W. Brown, C. Simpson, A. Barta, M. Kalyna, Transcriptome survey reveals increased complexity of the alternative splicing landscape in Arabidopsis. Genome Res 22, 1184–1195 (2012).

47. E. Remy et al., Intron retention in the 5’UTR of the novel ZIF2 transporter enhances translation to promote zinc tolerance in arabidopsis. PLoS Genet 10, e1004375 (2014).

48. S. Brogna, T. McLeod, M. Petric, The Meaning of NMD: Translate or Perish. Trends Genet 32, 395–407 (2016).

49. P. Akiva et al., Transcription-mediated gene fusion in the human genome. Genome Res 16, 30–36 (2006).

50. R. C. Allshire, H. D. Madhani, Ten principles of heterochromatin formation and function. Nat Rev Mol Cell Biol 19, 229–244 (2018).

51. Z. Zhong et al., DNA methylation-linked chromatin accessibility affects genomic architecture in Arabidopsis. Proc Natl Acad Sci U S A 118, (2021).

52. T. Preiss, M. Muckenthaler, M. W. Hentze, Poly(A)-tail-promoted translation in yeast: implications for translational control. RNA 4, 1321–1331 (1998).

53. Y. Zhang et al., The poly(A) polymerase PAPS1 interacts with the RNA-directed DNA-methylation pathway in sporophyte and pollen development. Plant J 99, 655–672 (2019).

54. S. San Paolo et al., Distinct roles of non-canonical poly(A) polymerases in RNA metabolism. PLoS Genet 5, e1000555 (2009).

55. H. Lange, F. M. Sement, J. Canaday, D. Gagliardi, Polyadenylation-assisted RNA degradation processes in plants. Trends Plant Sci 14, 497–504 (2009).

56. A. S. Reddy, Y. Marquez, M. Kalyna, A. Barta, Complexity of the alternative splicing landscape in plants. Plant Cell 25, 3657–3683 (2013).

57. R. K. Chodavarapu et al., Relationship between nucleosome positioning and DNA methylation. Nature 466, 388–392 (2010).

58. T. K. To, H. Saze, T. Kakutani, DNA Methylation within Transcribed Regions. Plant Physiol 168, 1219–1225 (2015).

59. A. Yearim et al., HP1 is involved in regulating the global impact of DNA methylation on alternative splicing. Cell Rep 10, 1122–1134 (2015).

60. B. Zhang et al., Linking key steps of microRNA biogenesis by TREX-2 and the nuclear pore complex in Arabidopsis. Nat Plants 6, 957–969 (2020).

61. R. R. Wick, L. M. Judd, K. E. Holt, Performance of neural network basecalling tools for Oxford Nanopore sequencing. Genome Biol 20, 129 (2019).

62. R. E. Workman et al., Nanopore native RNA sequencing of a human poly(A) transcriptome. Nat Methods 16, 1297–1305 (2019).

63. R. Li et al., Direct full-length RNA sequencing reveals unexpected transcriptome complexity during Caenorhabditis elegans development. Genome Res 30, 287–298 (2020).

64. M. D. Robinson, D. J. McCarthy, G. K. Smyth, edgeR: a Bioconductor package for differential expression analysis of digital gene expression data. Bioinformatics 26, 139–140 (2010).

65. F. Krueger, S. R. Andrews, Bismark: a flexible aligner and methylation caller for Bisulfite-Seq applications. Bioinformatics 27, 1571–1572 (2011).

66. A. R. Quinlan, I. M. Hall, BEDTools: a flexible suite of utilities for comparing genomic features. Bioinformatics 26, 841–842 (2010).

