## Supplemental Data 1 for "DNA methylation affects pre-mRNA transcriptional initiation and processing in Arabidopsis"

### Supplementary Information

**Additional file 1: Table S1**

Basic statistics of ONT DRS reads for Col-0, *met1-3*, *ddcc* and *tcx5tcx6*.

| | col-0 | met1-3 | ddcc | tcx5tcx6 |
|---|---|---|---|---|
| read* number | 2,336,056 | 2,204,601 | 2,583,621 | 2,302,562 |
| Average length | 963 | 1,057 | 1,008 | 875 |
| Median length | 826 | 899 | 859 | 739 |
| Average read quality | 10 | 10 | 10 | 10 |
| Alignment identity | 89.9% | 89.9% | 89.3% | 89.0% |
| Reads with poly-A (detected by nanopolish) | 2,206,865 (94.5%) | 2,104,047 (95.4%) | 2,470,373 (95.6%) | 2,165,155 (94.0%) |

*reads are high-quality (Qscore ≥7)

**Table S2:** Primers used in this study

| | |
|---|---|
| at4g04810-F | TTCTTTCTCCTGAACAGTTTCG |
| at4g04830-R1 | AGGAGTCTTGCATCCTACG |
| at4g04830-R2 | ATCGGTTGGATTACCGTAACC |
| at5g56010-F | AGACAAGAGCAAGCTCGATGG |
| at5g56030-R1 | AACTTCCTCAACCTTGCCTTCC |
| at5g56030-R2 | AACAACAACCTTGTCAGCAACC |
| at2g22670(met1-3)-F | ATCAATCGTTTCTCGACCTCC |
| at2g22670(met1-3)-R1 | AACACCAAGACCAGGTTTCC |
| at2g22670(met1-3)-R2 | TTGTCTTTAGAAGGTAGCAACG |
| new-over-at2g11410-420-F | AACTGAGATCGTGGTCTCG |
| new-over-at2g11410-420-R1 | TACATGAAGACGCTCTTCG |
| new-over-at2g11410-420-R2 | AACGTAGCTTAGTGATGTCAAGC |
| ACT7-F | ATCAATCCTTGCATCCCTCAG |
| ACT7-R | ACCACGAACCAGATAAGACAAG |
| At1g02750- For intron retention-F | ATCGATCGGAAACTGCTGC |
| At1g02750- For intron retention-R | TGGCAACTCATCAAGATGAGG |
| at3g06620-4th intron-retention-F | TACTGATAGCGAAGGCTTGG |
| at3g06620-4th intron-retention-R | AACAGCAGGACGTTAGGGTG |
| AT1G28060-F | ATGGCGGAAACAAGTGTTGG |
| AT1G28070-R | AGAAGTGAGAACTGAGCTCC |
| at4g15950-F | TTGGAGGAGCTATCCAAGC |
| at4g15953-R | ATCCGGTTAACCTCCTCG |
| at4g03157-F | AAGGTTTGATGATGAGGATGTGG |
| at4g03156-R | ATCTGACCATCTTCCTCAGC |
| at3g60400-F | TTGTGGACCTGAACACAAAGG |
| at3g60410-R | ATCGGAACCATGGATCGGTGG |
| at4g00030-F | AACTGGGAGCTCCCACTGC |
| at4g00040-R | TTGCTCGGCAAGGCCTTACC |

**Additional file 2: Figure S1-Figure S12.**

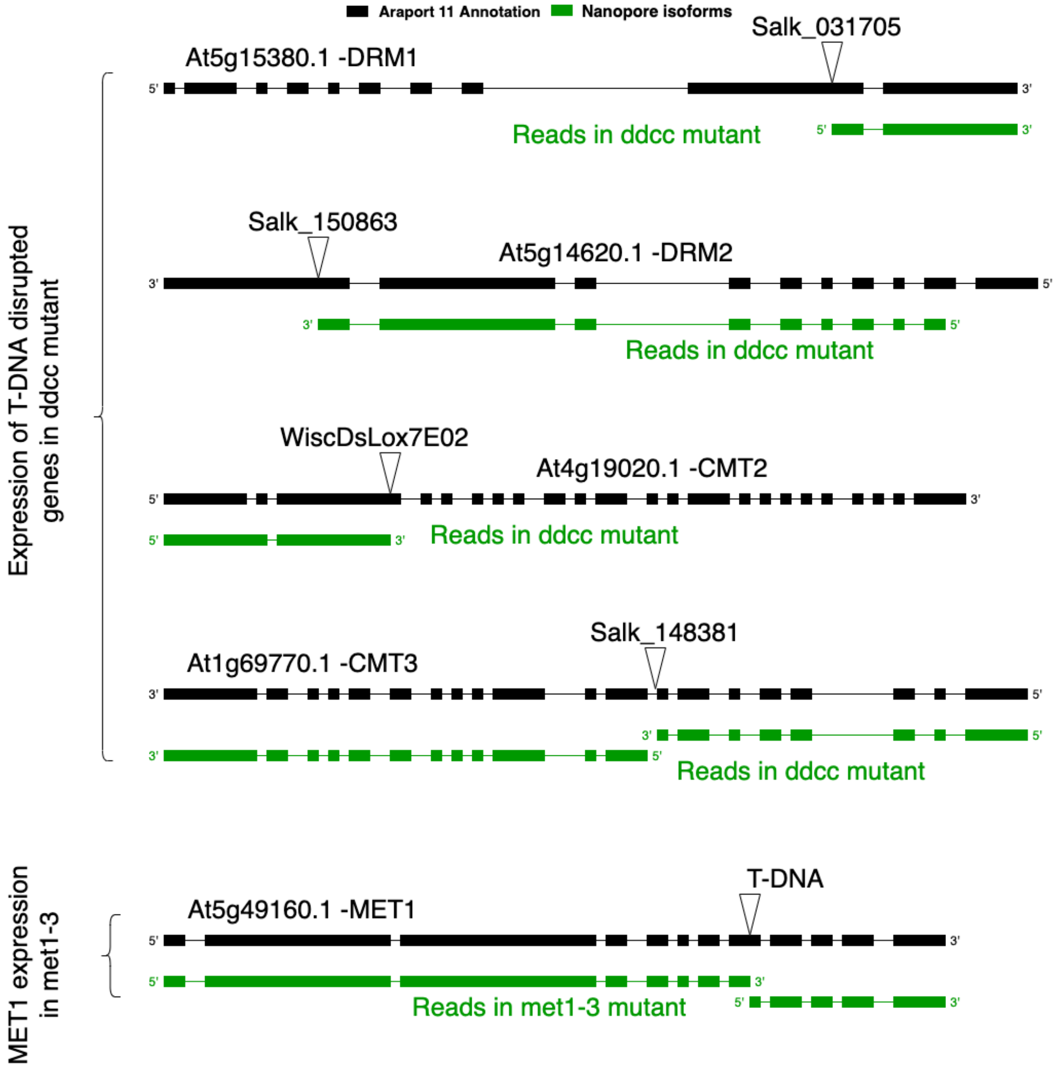


**Figure S1.** Aberrant transcripts from T-DNA insertion genes detected with ONT DRS reads (illustrated based on sequenced ONT DRS reads). Black box: exon annotated in Araport11; black line: intron annotated in Araport11; green box: exon detected with ONT DRS; green line: putative intron annotated according to ONT DRS mapping data; upside-down triangle: T-DNA inserted sites**.**

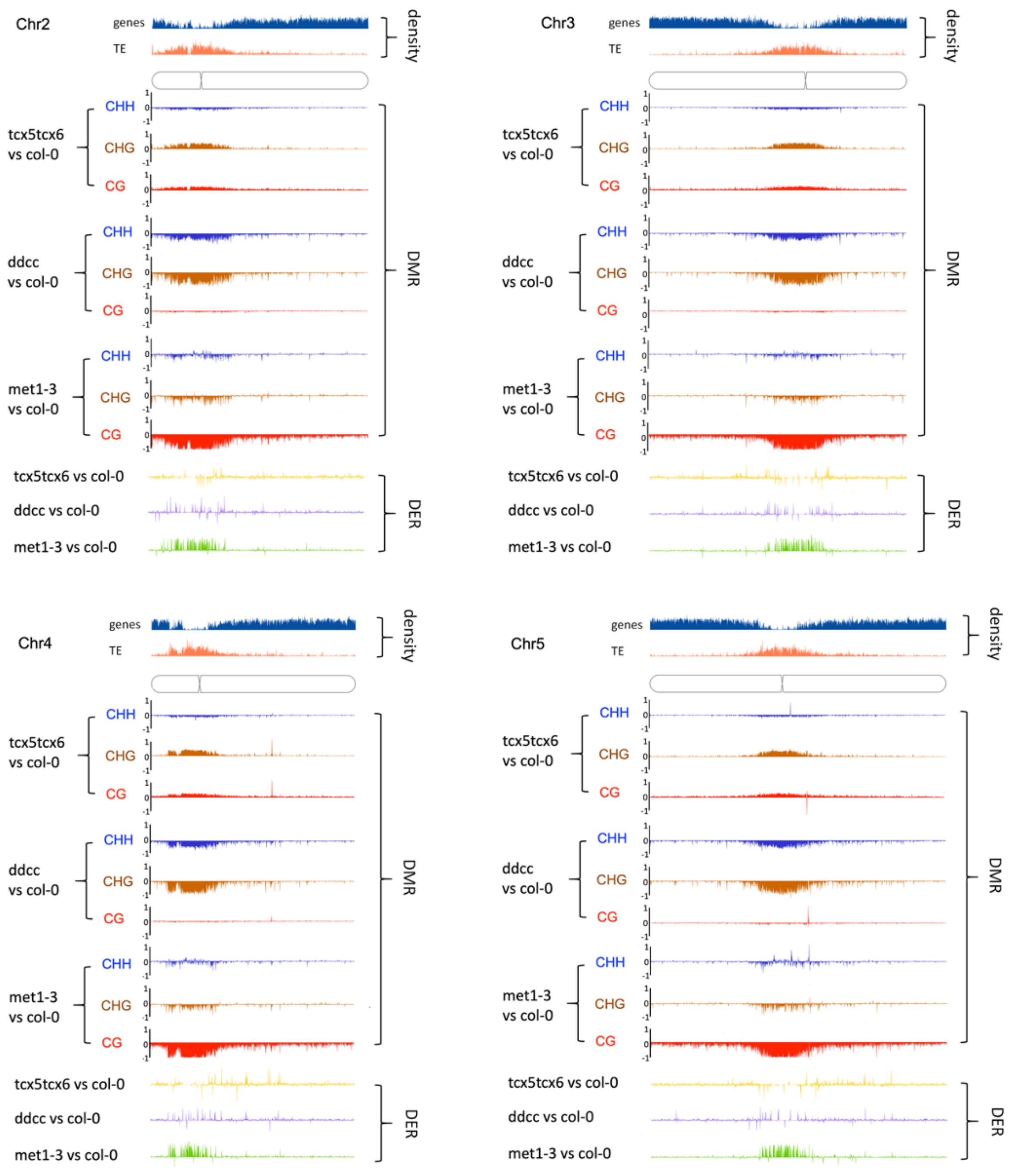


**Figure S2.** The differential expression regions (DER) and differential methylation regions (DMR) along various chromosomes between DNA methylation-related mutants (*met1-3/ddcc/tcx5tcx6*) and wild type (Col-0).

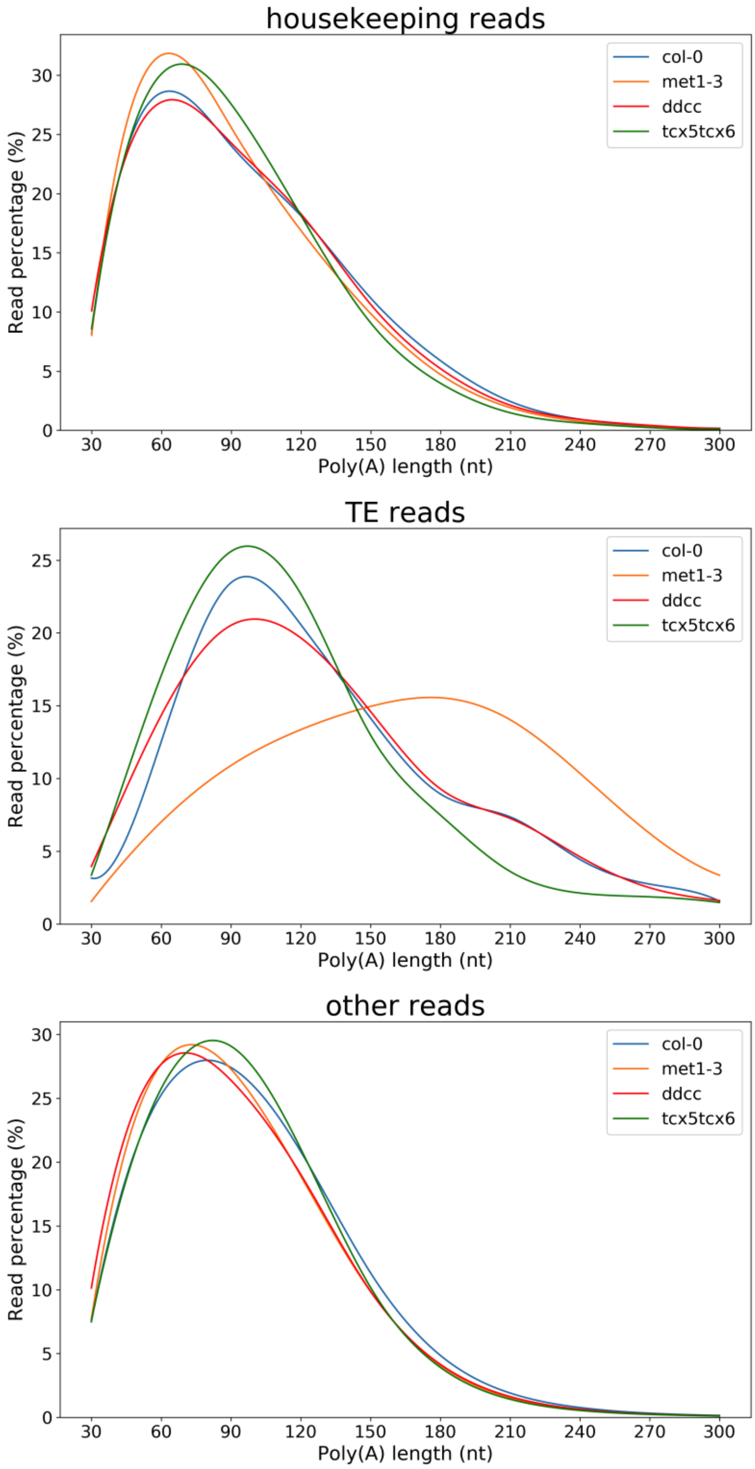


**Figure S3.** Comparison of poly (A) tail lengths among four genotypes for housekeeping genes (upper panel), transposable element (TE) genes (middle panel), and other genes (lower panel).

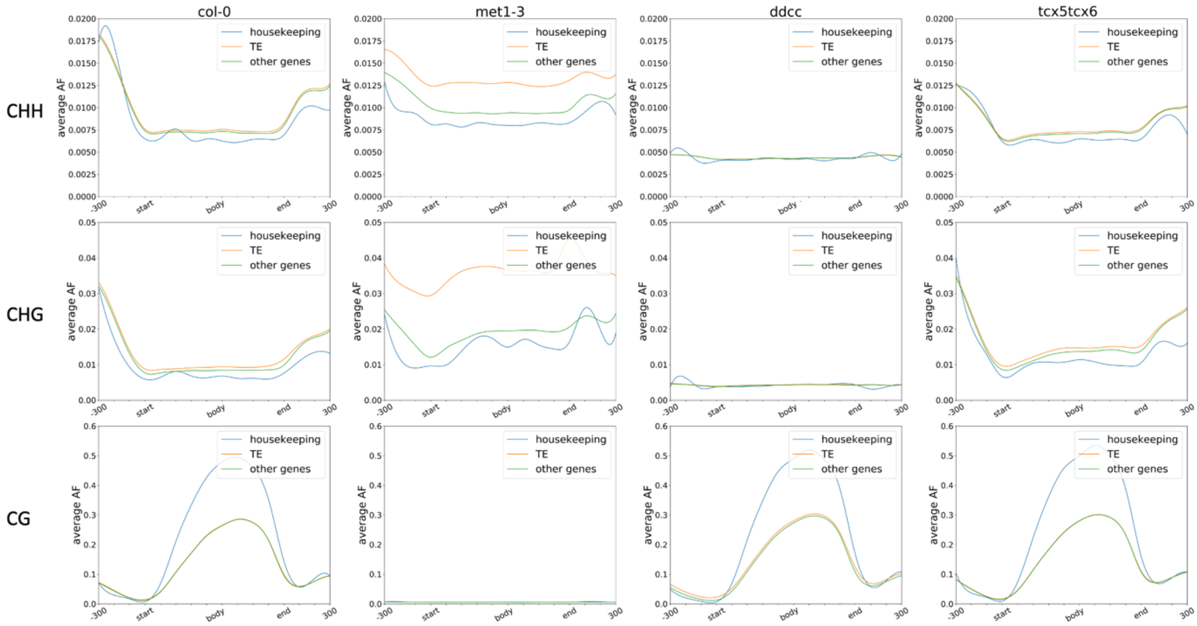


**Figure S4.** DNA methylation patterns for housekeeping genes, TE genes, and other genes in each genotype in the CHH, CHG, and CG contexts.

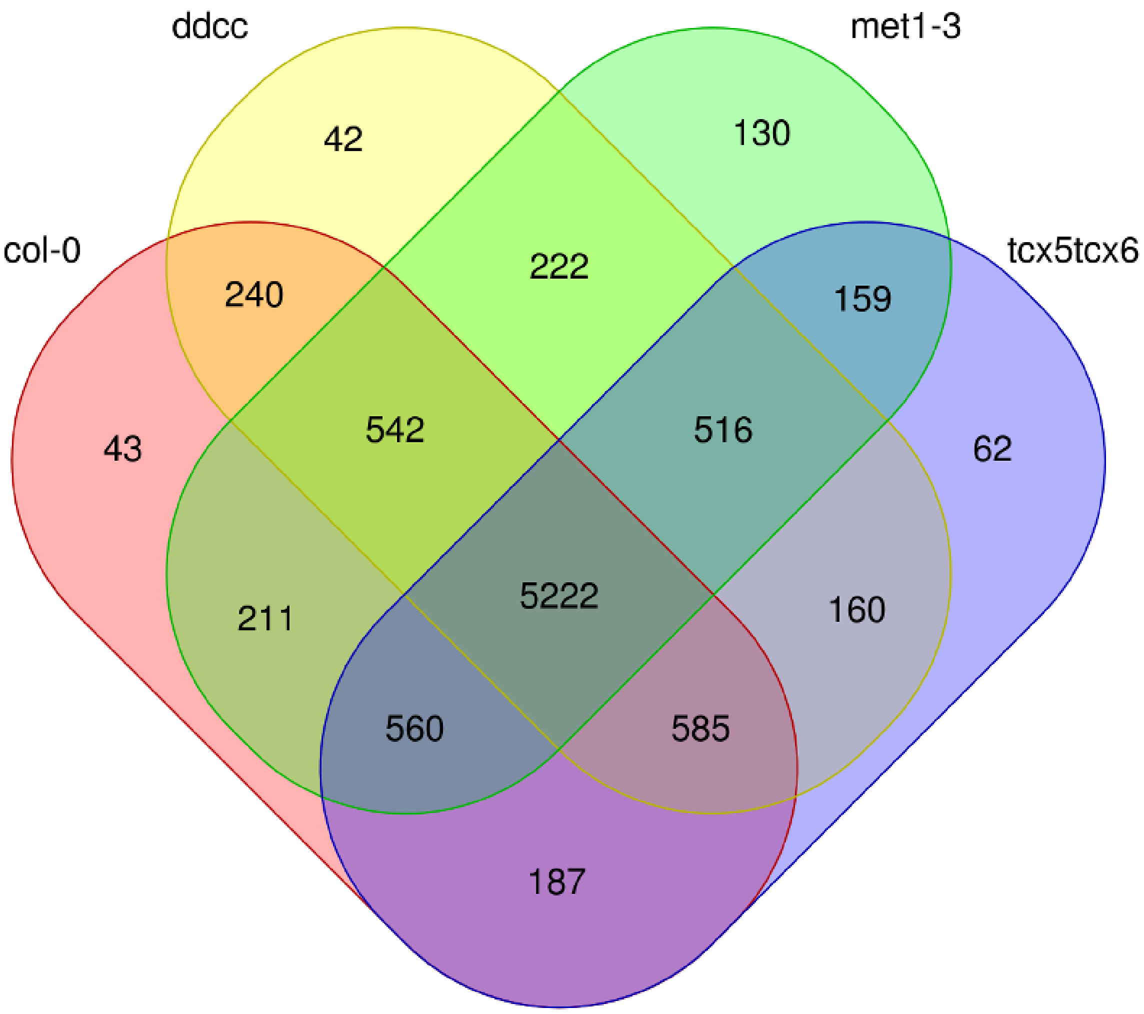


**Figure S5.** Venn diagram showing the number of intron retention events occurring in each genotype.

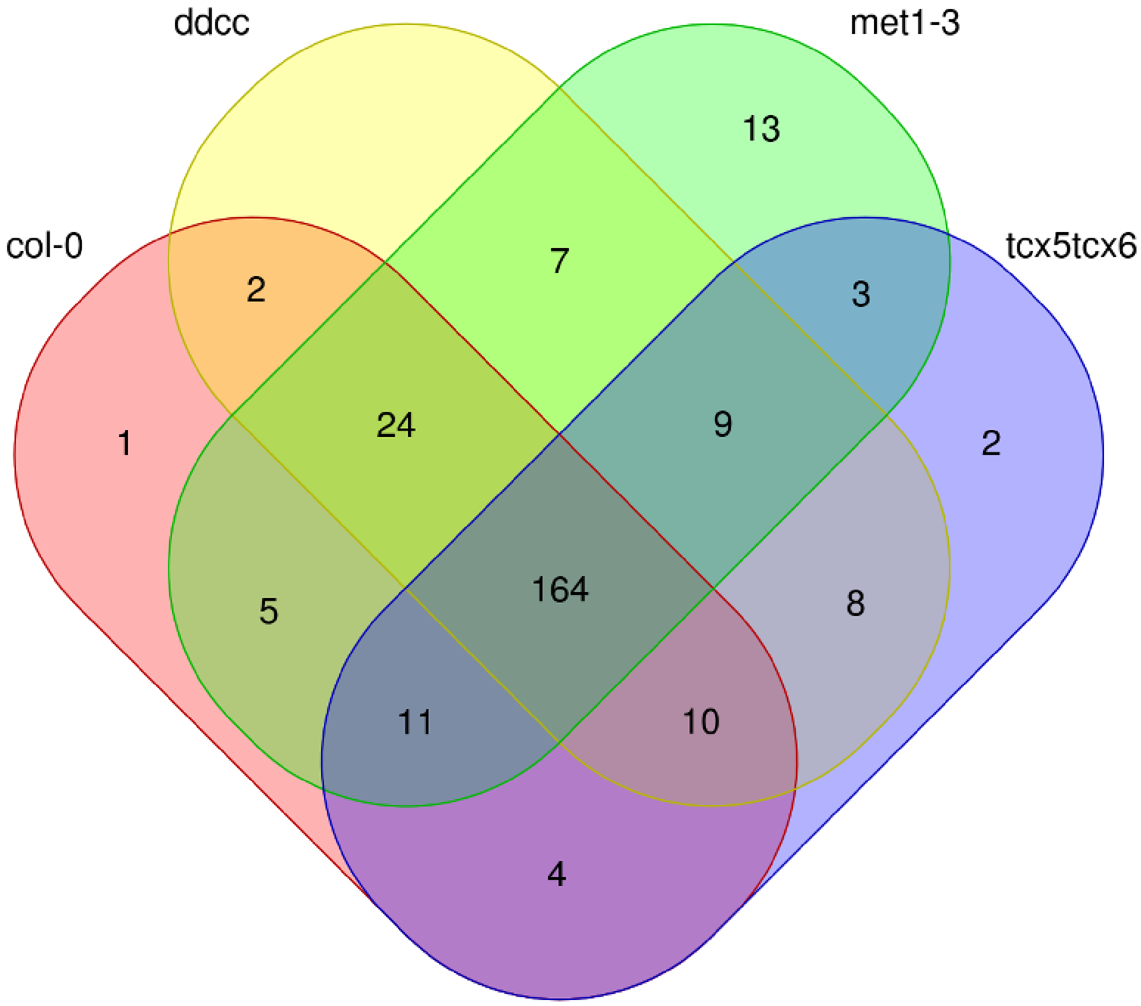


**Figure S6.** Venn diagram showing the number of extra exon events occurring in each genotype.

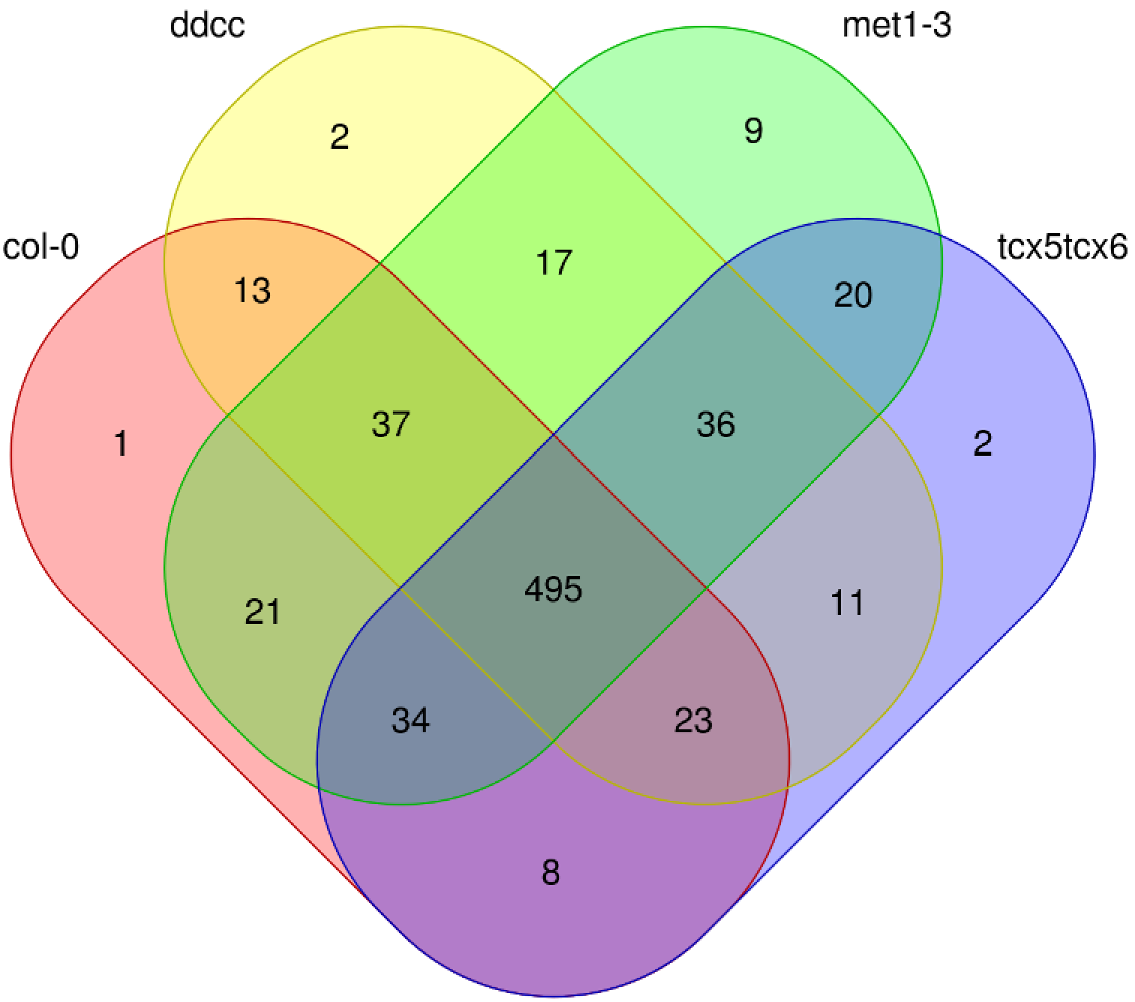


**Figure S7.** Venn diagram showing the number of exon-skipping events occurring in each genotype.

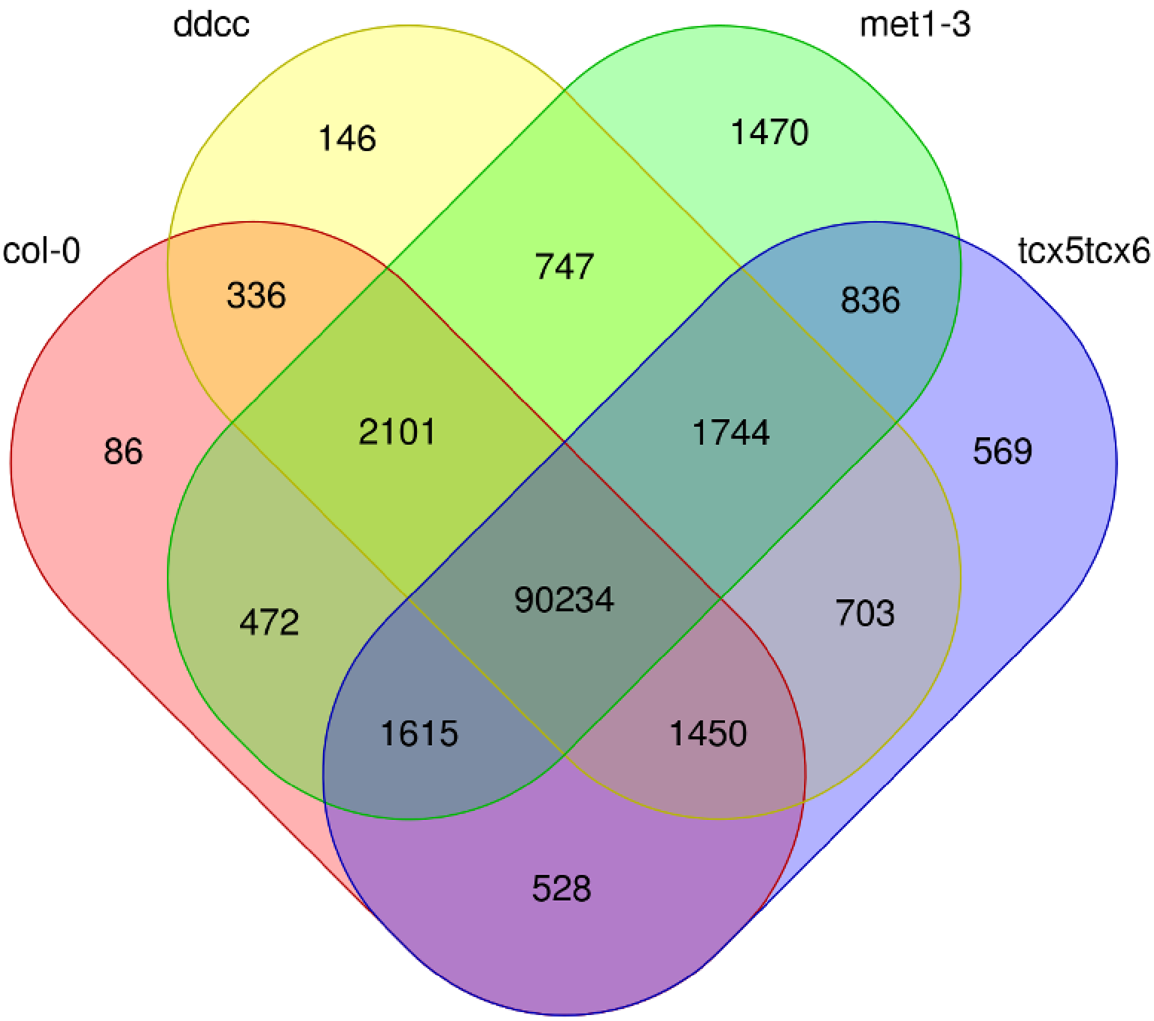


**Figure S8.** Venn diagram showing the number of identified splicing acceptor sites in the four genotypes.

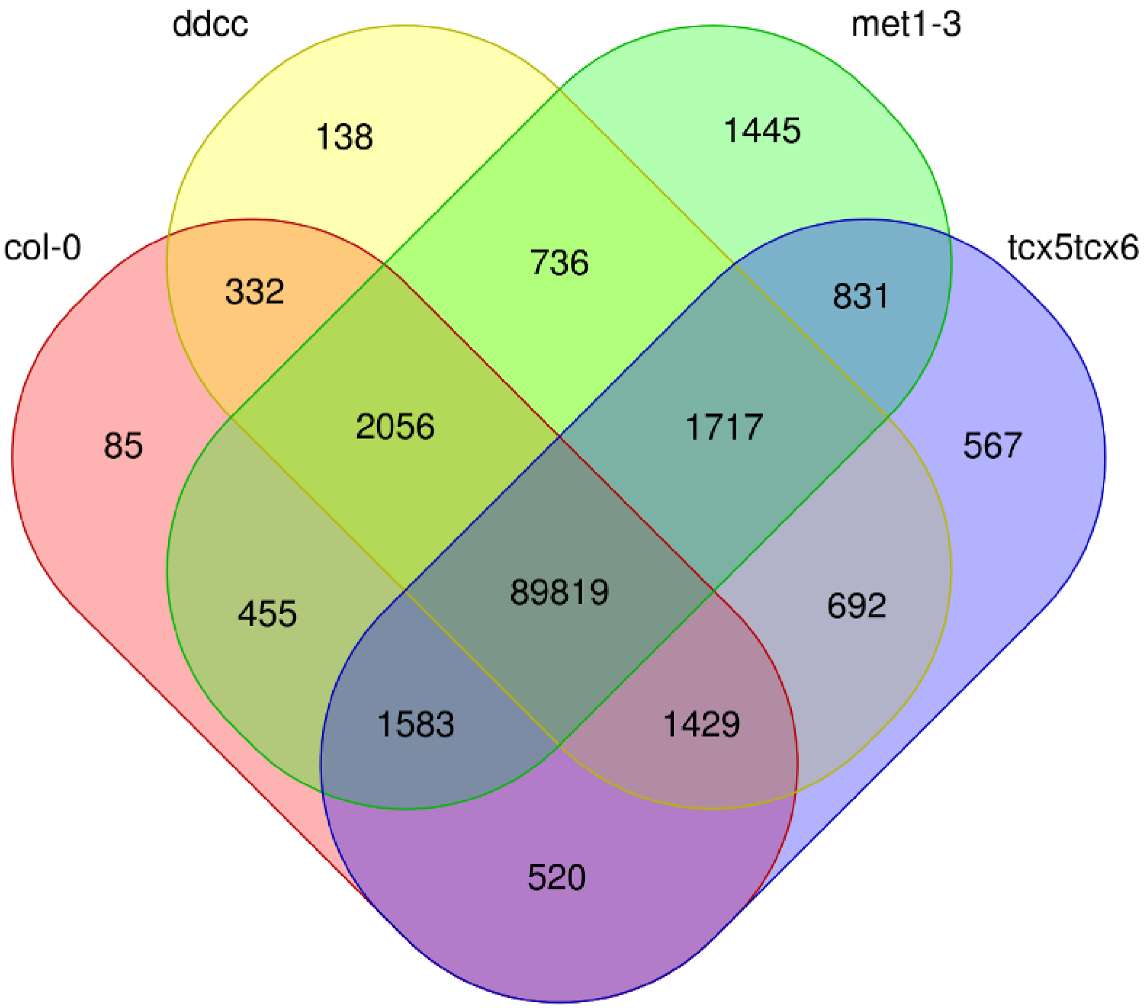


**Figure S9.** Venn diagram showing the number of identified splicing donor sites in the four genotypes.

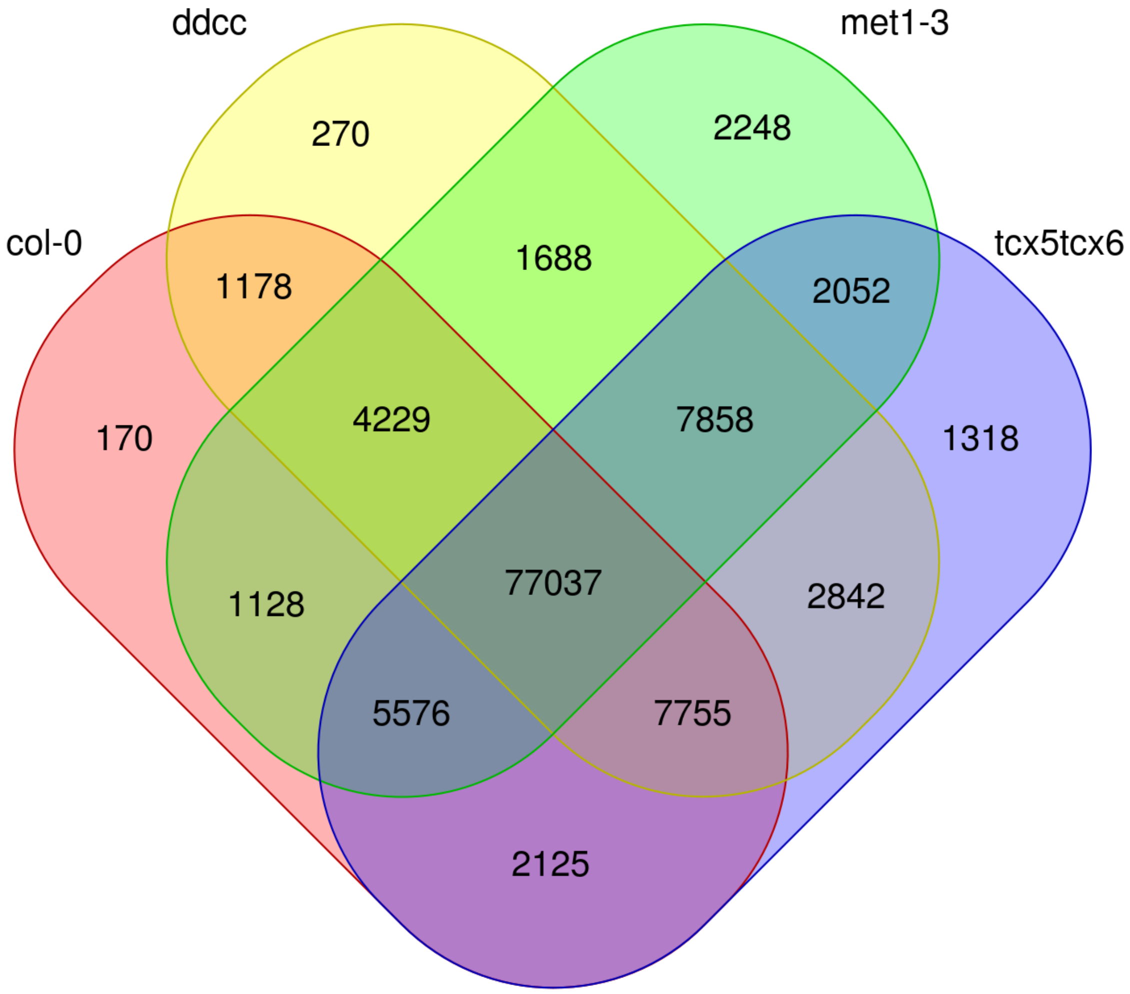


**Figure S10.** Venn diagram indicating the number of TC-RENO-identified transcript isoforms among the four genotypes.

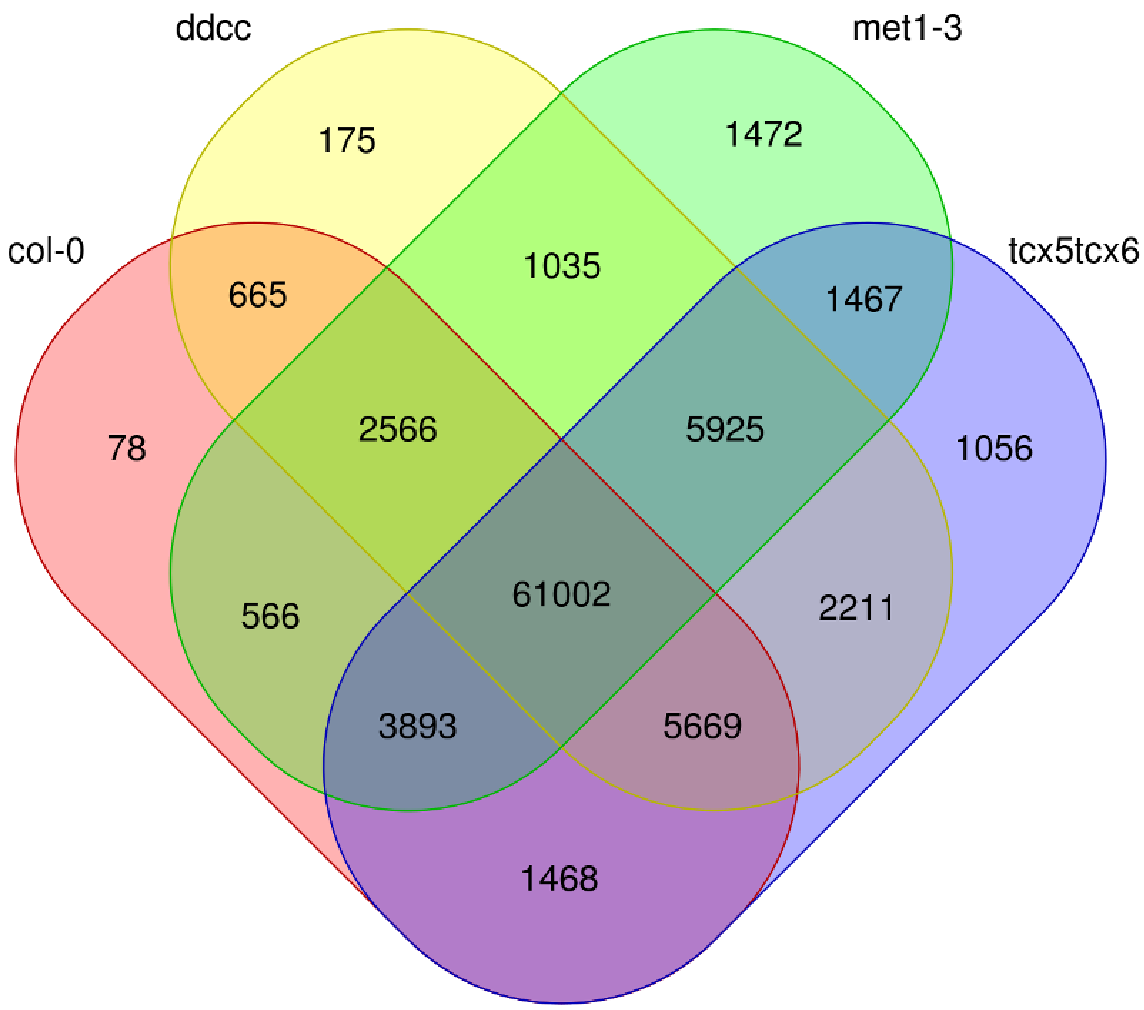


**Figure S11.** Venn diagram showing the numbers of transcriptional start sites (TSSs) in each genotype and the number of TSSs in common between genotypes.

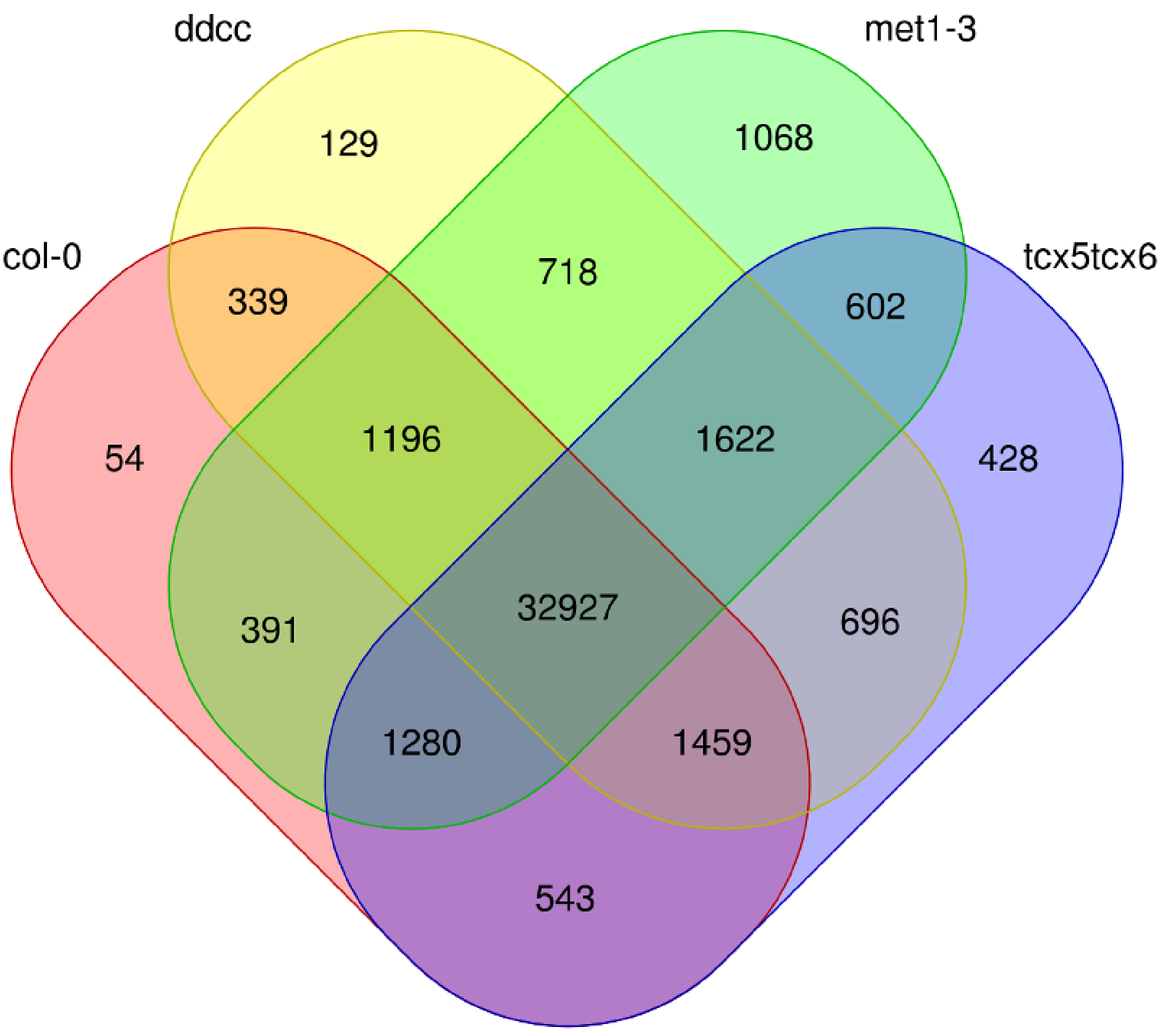


**Figure S12.** Venn diagram showing the numbers of transcriptional termination sites (TTSs) in each genotype and the number of TTSs in common between genotypes.

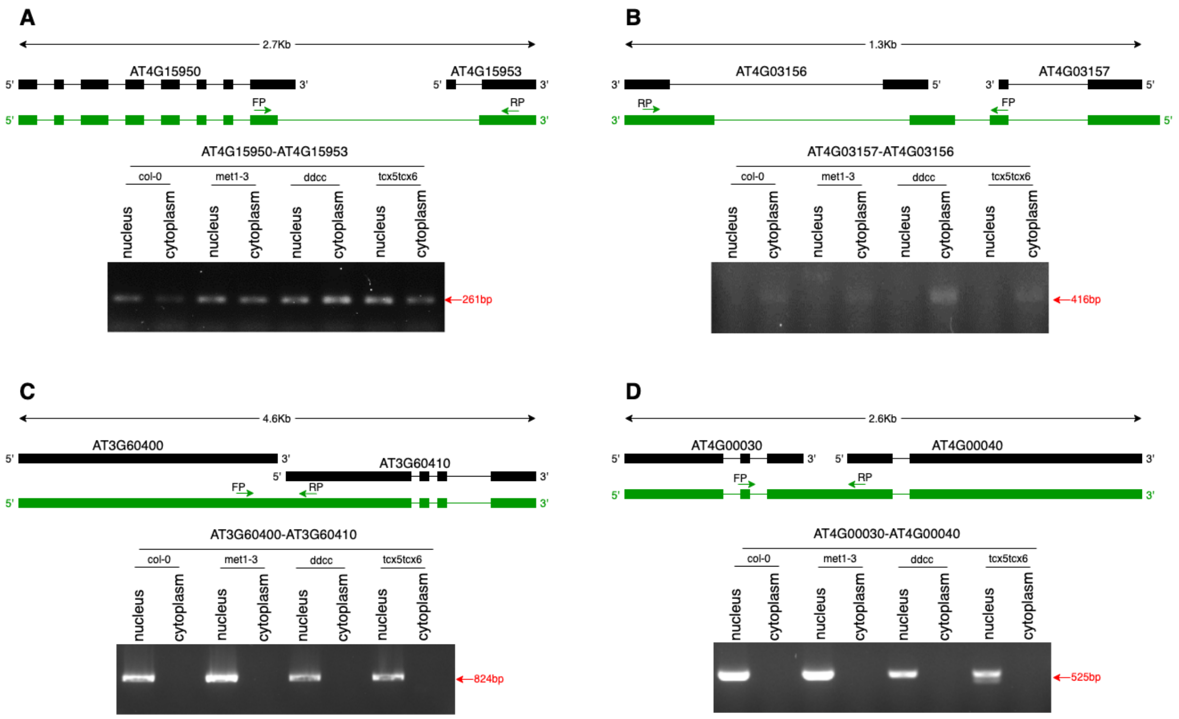


**Figure S13.** Detection of Arabidopsis fusion transcripts in the nucleus and the cytoplasm by RT-PCR and visualized by chemiluminescence detection. A-B. Two representative fusion transcripts without sequences transcribed from intergenic regions were able to be exported to the cytoplasm. C-D. Two representative fusion transcripts with sequences transcribed from intergenic regions were both retained in the nucleus. met1-3/ddcc/tcx5tcx6: DNA methylation-related mutants; col-0: wild type; black box: exon annotated in Araport11; black line: intron annotated in Araport11; green box: exon detected with ONT DRS; green line: putative intron annotated according to ONT DRS mapping data; FP: forward primer; RP: reverse primer. Green arrow indicates the primer orientation.

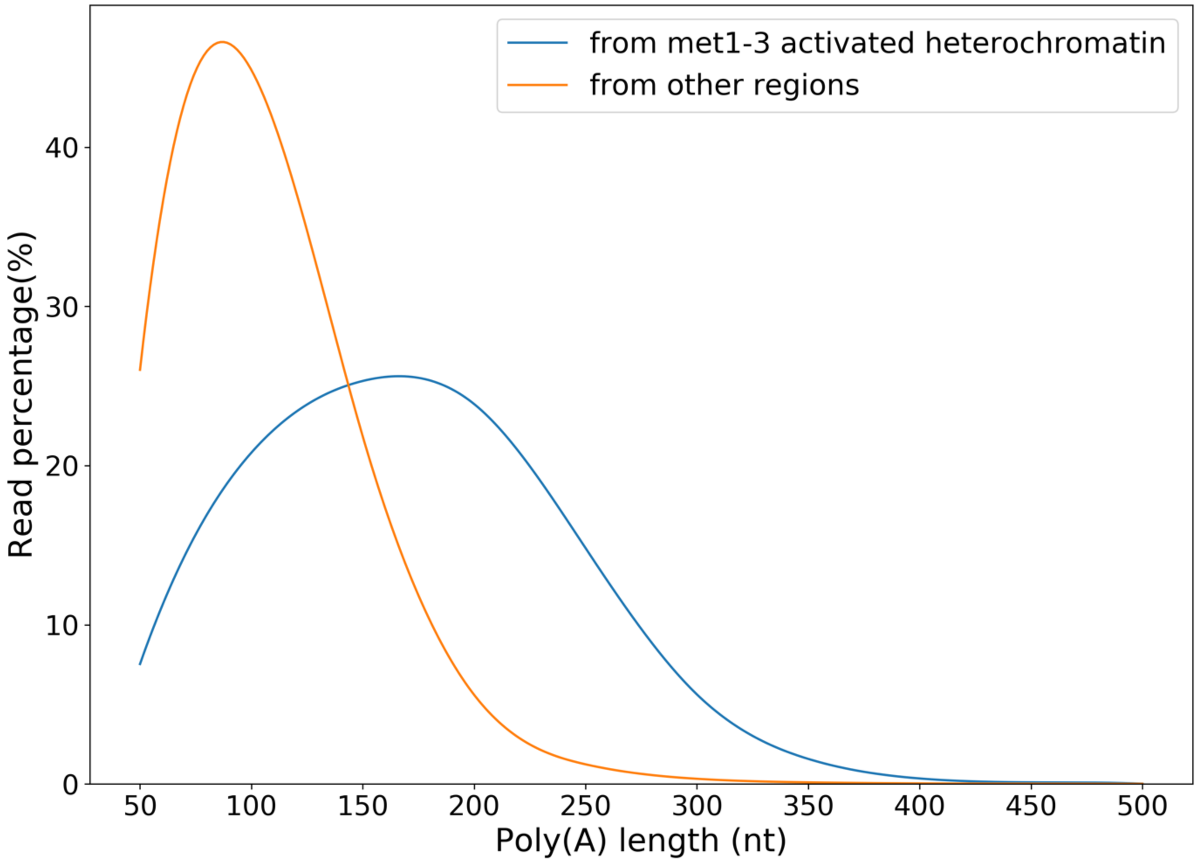


**Figure S14.** Comparison of the poly(A) tail lengths between euchromatin-transcribed transcripts and the constitutive heterochromatin-transcribed transcripts uniquely found in *met1-3*.
